## Appendix S6 for "Imaging with spatio-temporal modelling to characterize the dynamics of plant-pathogen lesions"

### S6 Comparison of estimated parameters

#### S6.1 Estimated parameters

| | Cultivar | Leaf Surface ($cm^2$) | Diffusion $\hat{D}$ | Growth rate $\hat{a}$ | Cost function J |
|---|---|---|---|---|---|
| 1 | Solara | 7.59 | 4.51e-01 | 4.84e-01 | 1.76e-02 |
| 2 | Solara | 6.17 | 5.49e-01 | 5.53e-01 | 1.99e-02 |
| 3 | Solara | 7.41 | 4.98e-01 | 5.10e-01 | 1.65e-02 |
| 4 | Solara | 5.14 | 5.07e-01 | 5.24e-01 | 1.73e-02 |
| 5 | Solara | 7.24 | 5.34e-01 | 6.88e-01 | 1.39e-02 |
| 6 | Solara | 7.10 | 5.66e-01 | 5.20e-01 | 1.93e-02 |
| 7 | Solara | 7.33 | 4.88e-01 | 4.61e-01 | 1.72e-02 |
| 8 | Solara | 8.98 | 5.85e-01 | 4.99e-01 | 1.86e-02 |
| 9 | Solara | 10.10 | 4.76e-01 | 4.42e-01 | 1.52e-02 |
| 10 | Solara | 8.66 | 6.89e-01 | 6.02e-01 | 1.74e-02 |
| 11 | Solara | 8.57 | 6.84e-01 | 5.44e-01 | 1.95e-02 |
| 12 | Solara | 9.32 | 5.14e-01 | 5.58e-01 | 1.54e-02 |
| 13 | Solara | 8.26 | 3.99e-01 | 6.37e-01 | 1.42e-02 |
| 14 | Solara | 8.40 | 3.04e-01 | 4.88e-01 | 1.51e-02 |
| 15 | Solara | 9.99 | 1.94e-01 | 5.42e-01 | 1.25e-02 |
| 16 | Solara | 6.60 | 3.78e-01 | 5.23e-01 | 1.52e-02 |
| 17 | James | 5.21 | 2.37e-01 | 4.49e-01 | 1.01e-02 |
| 18 | James | 3.90 | 1.83e-01 | 6.21e-01 | 8.51e-03 |
| 19 | James | 4.77 | 3.33e-01 | 5.16e-01 | 1.11e-02 |
| 20 | James | 4.99 | 3.19e-01 | 4.52e-01 | 1.22e-02 |
| 21 | James | 5.63 | 2.70e-01 | 4.44e-01 | 9.66e-03 |
| 22 | James | 4.83 | 2.93e-01 | 4.35e-01 | 1.14e-02 |
| 23 | James | 4.18 | 3.75e-01 | 4.65e-01 | 1.22e-02 |
| 24 | James | 4.32 | 2.60e-01 | 5.08e-01 | 1.22e-02 |
| 25 | James | 5.16 | 2.96e-01 | 4.73e-01 | 1.15e-02 |
| 26 | James | 4.54 | 4.59e-01 | 5.22e-01 | 1.51e-02 |
| 27 | James | 2.80 | 6.69e-02 | 4.16e-01 | 7.96e-03 |
| 28 | James | 4.78 | 4.08e-01 | 5.30e-01 | 1.37e-02 |
| 29 | James | 4.28 | 2.00e-01 | 6.04e-01 | 9.92e-03 |
| 30 | James | 4.00 | 2.87e-01 | 4.88e-01 | 1.27e-02 |
| 31 | James | 4.02 | 3.19e-01 | 5.00e-01 | 1.06e-02 |
| 32 | James | 4.39 | 3.55e-01 | 5.09e-01 | 1.15e-02 |

Table A: Estimated diffusion $\hat{D}$, rate growth $\hat{a}$ and cost function $J$ for 32 different sets of images.

### S6.2 Cultivar effect on estimated parameters

| | | Df | Sum Sq | Mean Sq | F value | Pr(>F) |
|---|---|---|---|---|---|---|
| Diffusion $\hat{D}$ | | | | | | |
| | Cultivar | 1 | 0.31 | 0.31 | 24.95 | 2.36e-05 |
| | Residuals | 30 | 0.37 | 0.01 | | |
| Growth rate $\hat{a}$ | | | | | | |
| | Cultivar | 1 | 0.01 | 0.01 | 3.56 | 6.90e-02 |
| | Residuals | 30 | 0.11 | 0.00 | | |

Table B: Anova table for the cultivar effect on the diffusion $\hat{D}$ and the local growth rate $\hat{a}$. Although the cultivar effect is significant on both $\hat{D}$ and $\hat{a}$ it explains 84% of the variance for the diffusion and only 9% for the growth rate.
