## Appendix S5 for "Imaging with spatio-temporal modelling to characterize the dynamics of plant-pathogen lesions"

### S5 Visual assessment of stipules deformation

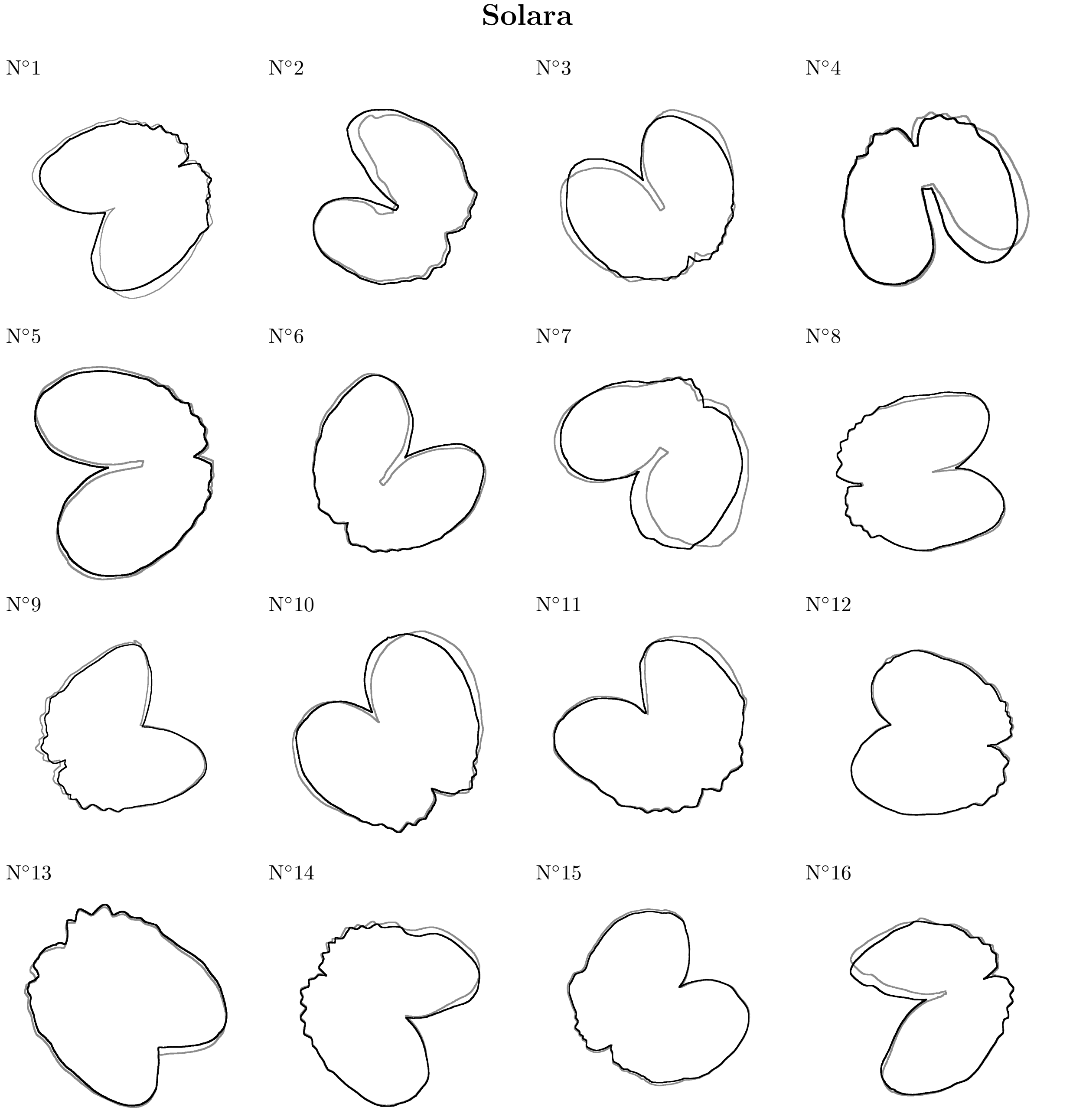


Figure A: Comparison of registered pea stipules for Solara. Stipules edges at day three are in black while those at day seven are in grey.

### James

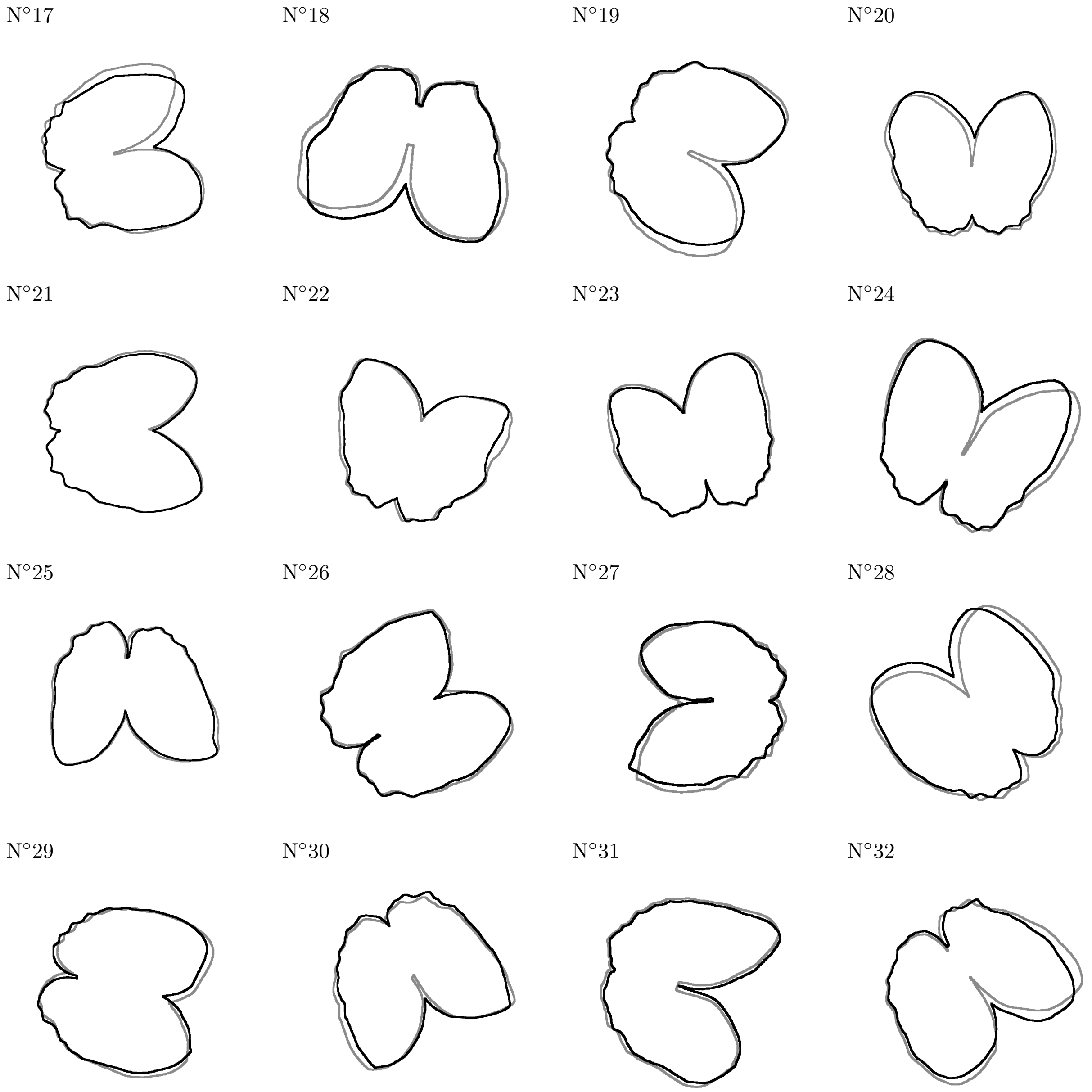


Figure B: Comparison of registered pea stipules for James. Stipules edges at day three are in black while those at day seven are in grey.
