## Appendix S4 for "Imaging with spatio-temporal modelling to characterize the dynamics of plant-pathogen lesions"

### S4 Numerical discretization and resolution of the optimization problem

#### S4.1 Meshing the computation domain

The level-set formalism is used to described the stipules surface and boundary. Let $\phi$ be the level-set function such that the stipules boundary is the zero level of $\phi$ defined as

$$\partial\Omega := \left\{\mathbf{x} \in \mathbb{R}^2;\ \phi(\mathbf{x}) = 0\right\}.$$

Then the stipules surface is $\Omega := \left\{\mathbf{x} \in \mathbb{R}^2;\ \phi(\mathbf{x}) < 0\right\}$, and its complementary $\Omega^c := \left\{\mathbf{x} \in \mathbb{R}^2;\ \phi(\mathbf{x}) > 0\right\}$. The function $\phi$ easily provides the exterior normal of leaf as $\vec{n} = \frac{\nabla\phi}{||\nabla\phi||}$. The computation domain consists in a cartesian grid of the overall area (see Figure A). The partial differential equations are solved using explicit Euler finite differences in time and second order centered finite differences in space. The gradient is solved using a descent method [4, 5]. The method is implemented with Petsc using Python [2].

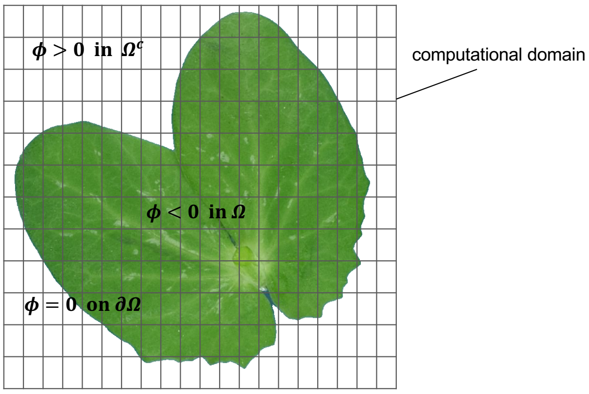


Figure A: Schematic representation of the computational domain given by a cartesian grid and the stipules area $\Omega = \left\{\mathbf{x} \in \mathbb{R}^2;\ \phi(\mathbf{x}) < 0\right\}$ defined by the level-set function $\phi$.

#### S4.2 Variational image data assimilation

The optimization problem is solved thanks to the minimization of the Lagrangian [1]

$$\begin{aligned}
\mathcal{L}(a, D) \quad &= \quad \frac{1}{2}\sum_{t=t_3}^{t_7}\int_{\Omega}(u(\mathbf{x},t,\theta) - u_{reg}(\mathbf{x},t))^2 d\mathbf{x} \\
&+ \sum_{t=t_3}^{t_7}\int_{\Omega}\bigg(\frac{\partial u(\mathbf{x},t,\theta)}{\partial t} - D\Delta u(\mathbf{x},t,\theta) \\
&\qquad - au(\mathbf{x},t,\theta)\left(1 - u(\mathbf{x},t,\theta)\right)\bigg)\lambda(\mathbf{x},t,\theta)d\mathbf{x}.
\end{aligned} \tag{Eq A}$$

The adjoint state $\lambda$ is the solution of the following backward PDE

$$-\frac{\partial \lambda}{\partial t} = D\Delta\lambda(\mathbf{x},t) + a\left(1 - 2u(\mathbf{x},t)\right)\lambda(\mathbf{x},t) + \left(u(\mathbf{x},t) - u_{reg}\right) \text{ in } \Omega \tag{Eq B}$$

with homogeneous Neummann boundary condition $\frac{\partial \lambda}{\partial n} = 0$ on $\partial\Omega$, and with the final condition $\lambda(t = t_7) = 0$. Finally, the gradient of the cost function is given by

$$\begin{cases}
\dfrac{\partial \mathcal{L}}{\partial D} = \displaystyle\sum_{t=t_3}^{t_7}\int_{\Omega}\nabla u(\mathbf{x},t)\nabla\lambda(\mathbf{x},t)d\mathbf{x} \\
\dfrac{\partial \mathcal{L}}{\partial a} = -\displaystyle\sum_{t=t_3}^{t_7}\int_{\Omega}u(\mathbf{x},t)\left(1 - u(\mathbf{x},t)\right)\lambda(\mathbf{x},t)d\mathbf{x}.
\end{cases} \tag{Eq C}$$

To avoid local minimum, the initial guess is determined by splitting the Fisher-KPP equation into

$$\frac{du}{dt}(\mathbf{x},t) \quad = \quad au(\mathbf{x},t)\left(1 - u(\mathbf{x},t)\right) \tag{Eq D}$$

$$\frac{\partial u}{\partial t}(\mathbf{x},t) \quad = \quad D\Delta u(\mathbf{x},t). \tag{Eq E}$$

From the given set of registerd images $u_{reg}$, the diffusion coefficient in equation (Eq D) is approximated with 98% of the population preservation [3, 6], by

$$D \simeq \frac{\langle r^2 \rangle}{16(t_b - t_a)}, \tag{Eq F}$$

where $\langle r^2 \rangle$ denotes the mean of the square radius of mycelium spread between times $t_b$ and $t_a$. On the other hand, the logistic equation (Eq E) has explicit solution written as

$$u(t_b) = \frac{1}{1 + \left(\frac{1}{u(t_a)} - 1\right)\exp(-a(t_b - t_a))}$$

that implies

$$a = -\frac{1}{(t_b - t_a)}\ln\left(\frac{\frac{1}{u(t_b)} - 1}{\frac{1}{u(t_a)} - 1}\right). \tag{Eq G}$$

The numerical procedure is summarized in Algorithm 1.

**Algorithm 1** Parameters identification

Given $N$ registered images $u_{reg}$ and a tolerance $\tau$
Compute the stipules area $\Omega$ using level-set
Compute initial $D_0$ with (Eq F) and $a_0$ with (Eq G)
**while** $||\nabla\mathcal{L}(\theta^m)|| \geq \tau$ **do**,
    Find $u^m(\mathbf{x}, t, \theta^m)$ solution of the forward PDE (1)
    Find $\lambda^m(\mathbf{x}, t, \theta^m, u^m)$ solution of the backward PDE (Eq B)
    Compute the gradient

$$\begin{cases} D^{m+1} = D^m - \rho_1^m \left( \displaystyle\sum_{t=t_3}^{t_7} \int_\Omega \nabla u^m(\mathbf{x}, t) \nabla \lambda^m(\mathbf{x}, t) d\mathbf{x} \right) \\ a^{m+1} = a^m + \rho_2^m \left( \sum t = {t_3}^{t_7} \displaystyle\int_\Omega u(\mathbf{x}, t) \left(1 - u(\mathbf{x}, t)\right) \lambda(\mathbf{x}, t) d\mathbf{x} \right) \end{cases}$$

#### S4.3 Numerical convergence examination

To ensure the convergence of the numerical method, the size of each set of images has been rescaled. Table A indicates that same order values are found for $a$. The diffusion coefficient depends on images size, but the relative diffusions, *i.e.* the quotient between diffusion coefficient and stipules area are equal at order $10^{-3}$. Convergence results are provided in Table A.

| | Cultivar | | | | | | |
|---|---|---|---|---|---|---|---|
| 1 | Solara | size | $512 \times 514$ | $614 \times 617$ | $717 \times 720$ | $819 \times 822$ | $922 \times 925$ |
| | | $\hat{a}$ | 0.4788 | 0.4835 | 0.4845 | 0.4852 | 0.4841 |
| | | $\hat{D}$ | 0.4597 | 0.4511 | 0.4529 | 0.4534 | 0.4584 |
| 17 | James | size | $512 \times 514$ | $614 \times 617$ | $717 \times 720$ | $819 \times 822$ | $922 \times 925$ |
| | | $\hat{a}$ | 0.4472 | 0.4492 | 0.4530 | 0.4550 | 0.4543 |
| | | $\hat{D}$ | 0.2367 | 0.2367 | 0.2368 | 0.2366 | 0.2368 |

Table A: Estimated diffusion $\hat{D}$, rate growth $\hat{a}$ for two different sets of images with five variable image sizes. The method provides same order parameters with respect to the size images.
