## Appendix S3 for "Imaging with spatio-temporal modelling to characterize the dynamics of plant-pathogen lesions"

- S3.1 Probability images, fitted models and images of residuals for all 32 inoculated stipules (p 2 - 33).
- S3.2 Boxplots showing the distributions of residuals for all dates and inoculated stipules (p 34 - 35).
- S3.3 Comparison of predicted infection probability against probability given by classifiers (pixel versus pixel) for all inoculated stipules (p 36 - 37).

#### S3 Assessment of the discrepancy between model and data

##### S3.1 Spatial comparison of model prediction against data

For each monitored inoculated stipules the discrepancy between the probability images (top) and the fitted reaction-diffusion model (middle) is assessed by visualizing the raw residuals $[u_{reg}(\mathbf{x}, t_i) - u(\mathbf{x}, t_i, \hat{\theta})]$ (bottom) at day 4, 5, 6 and 7 after inoculation (from left to right).

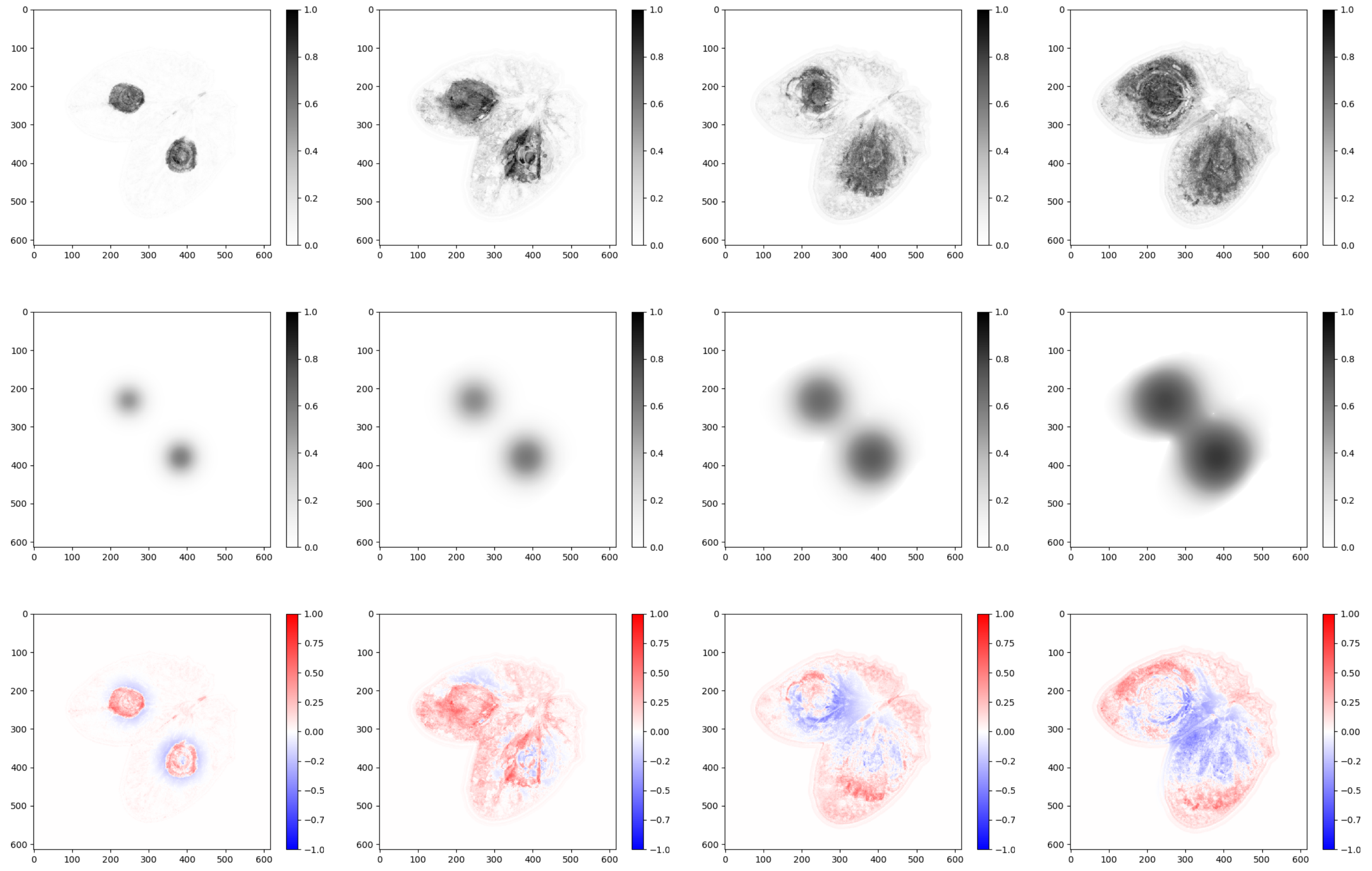

Figure A: Solara N°1

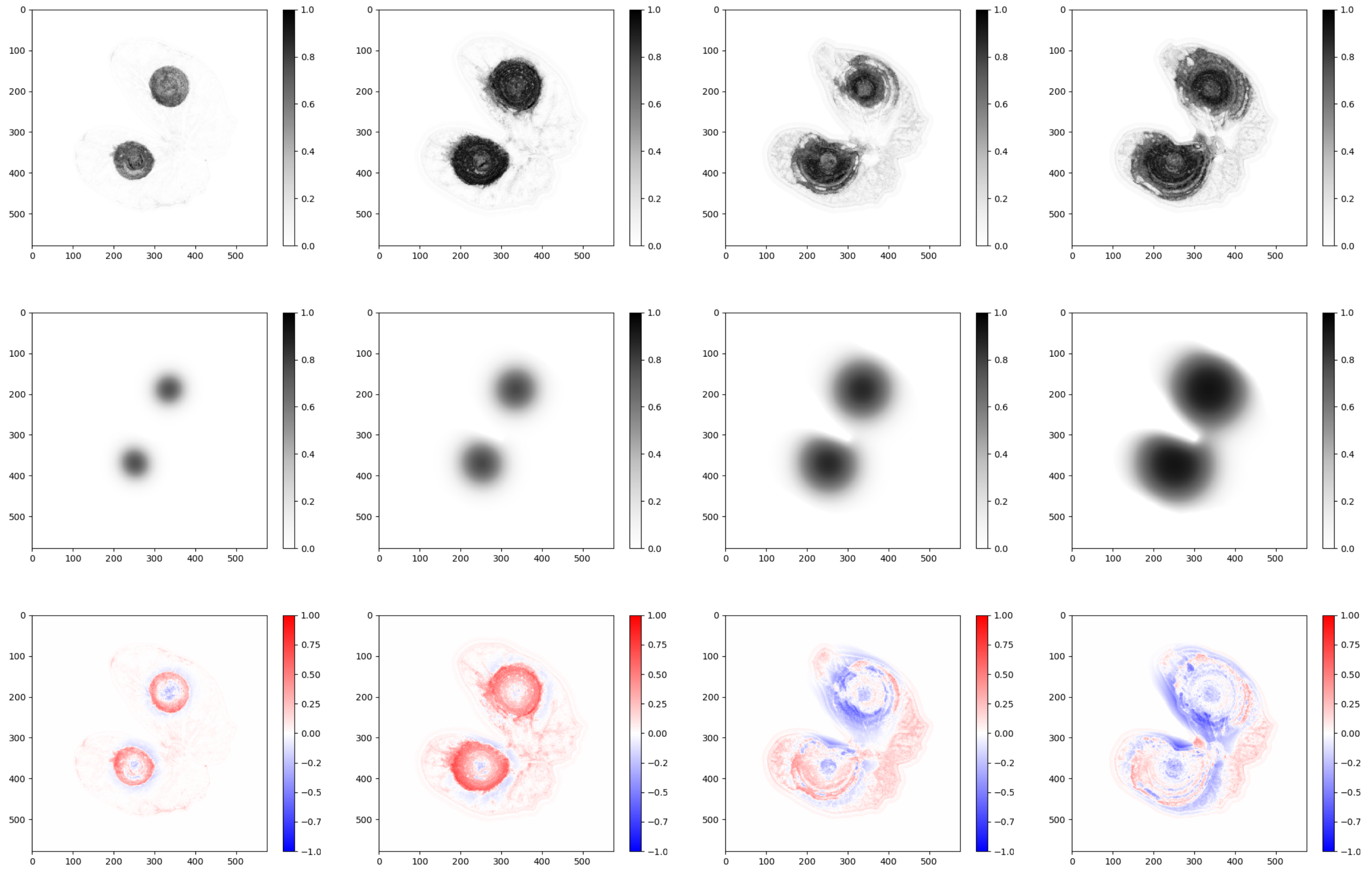


Figure B: Solara N°2

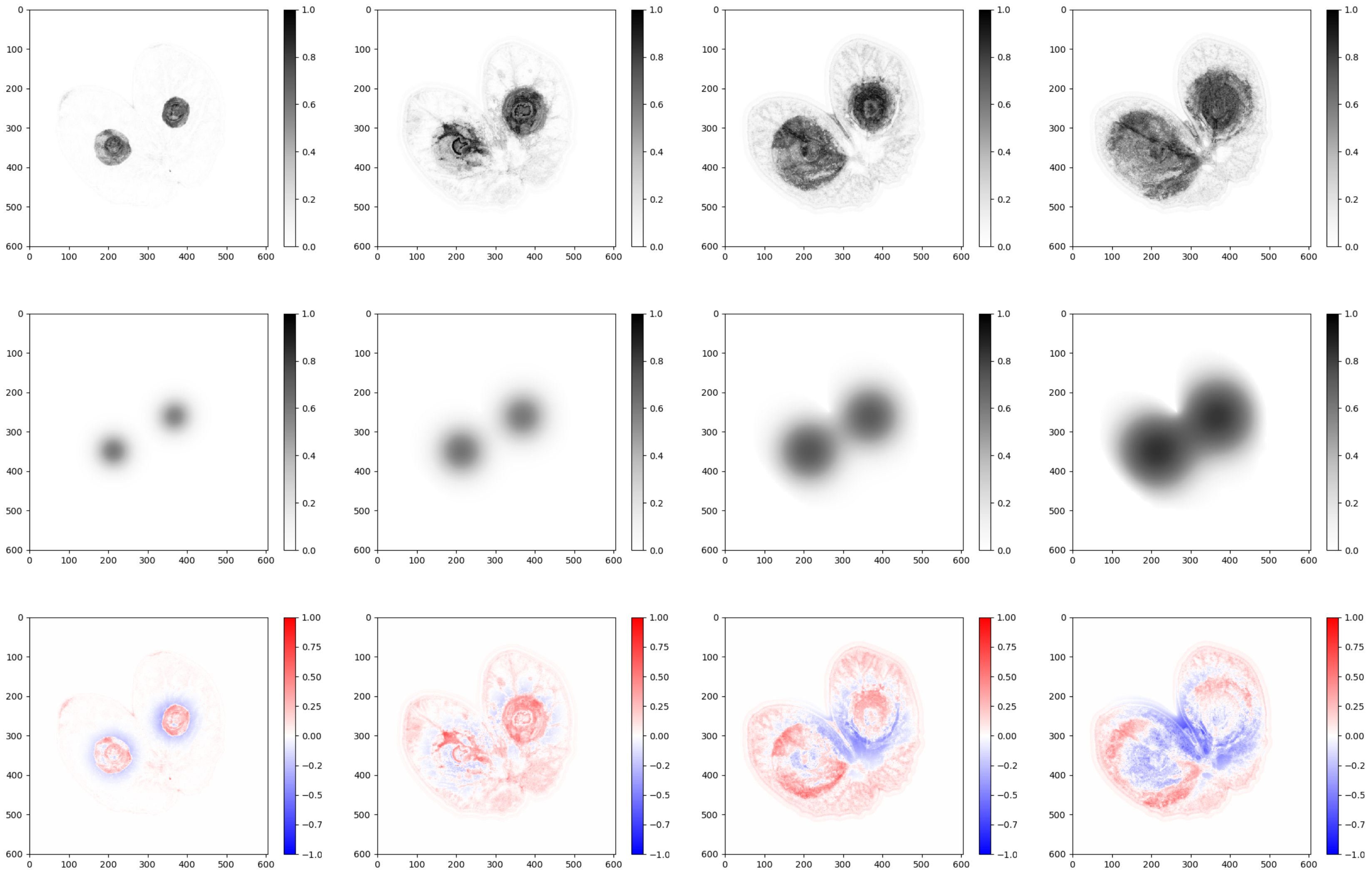

Figure C: Solara N°3

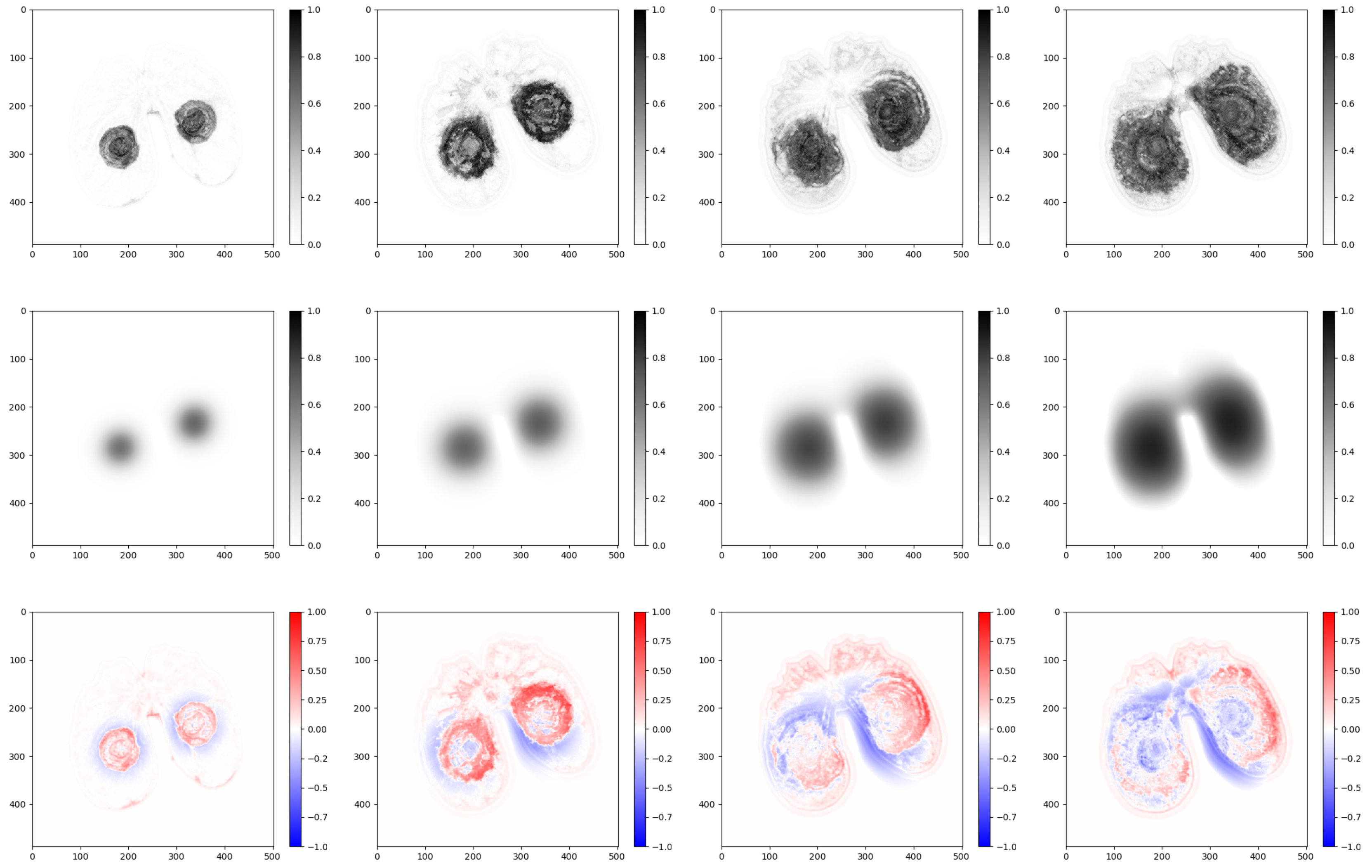


Figure D: Solara N°4

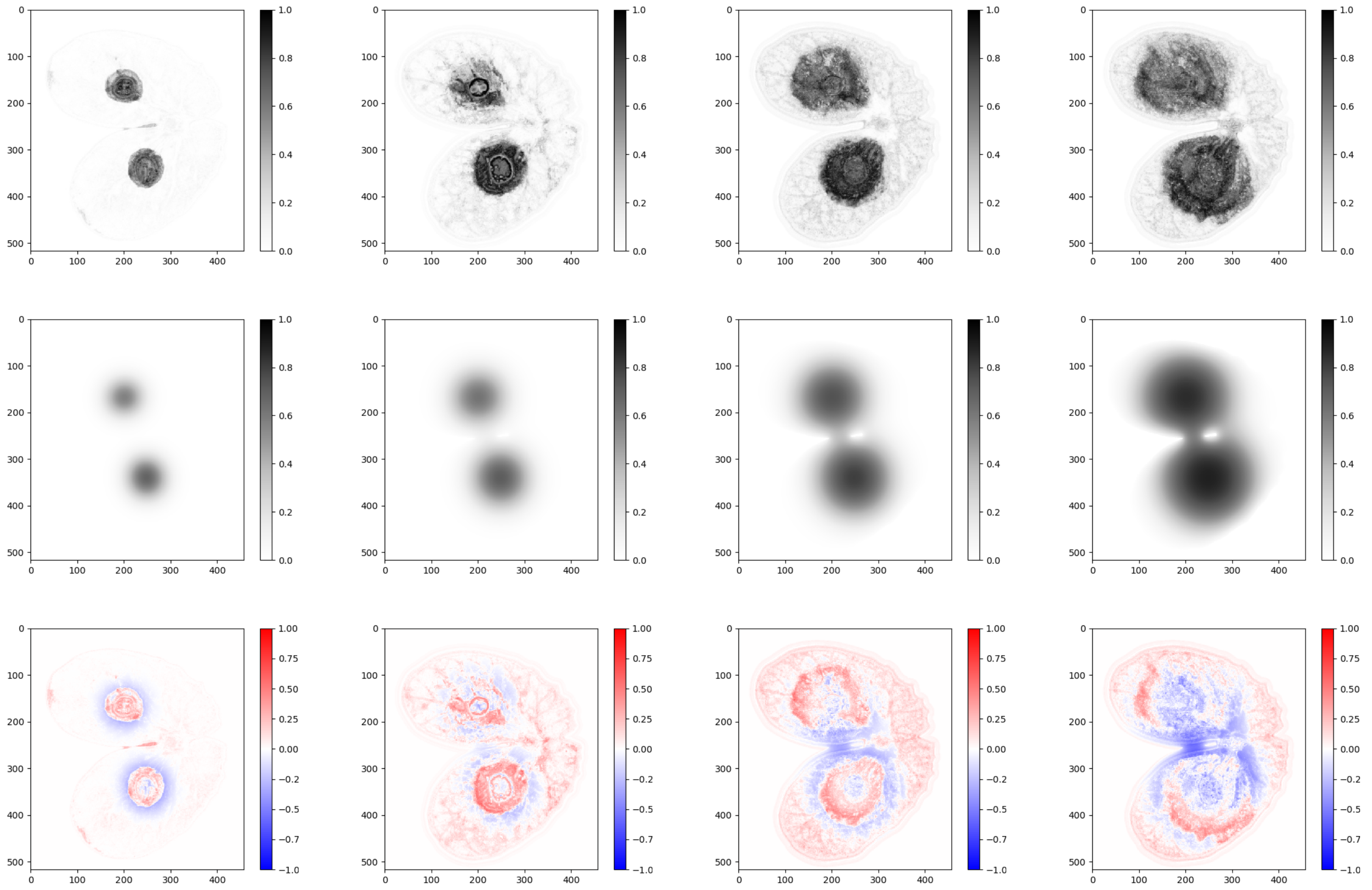


Figure E: Solara N°5

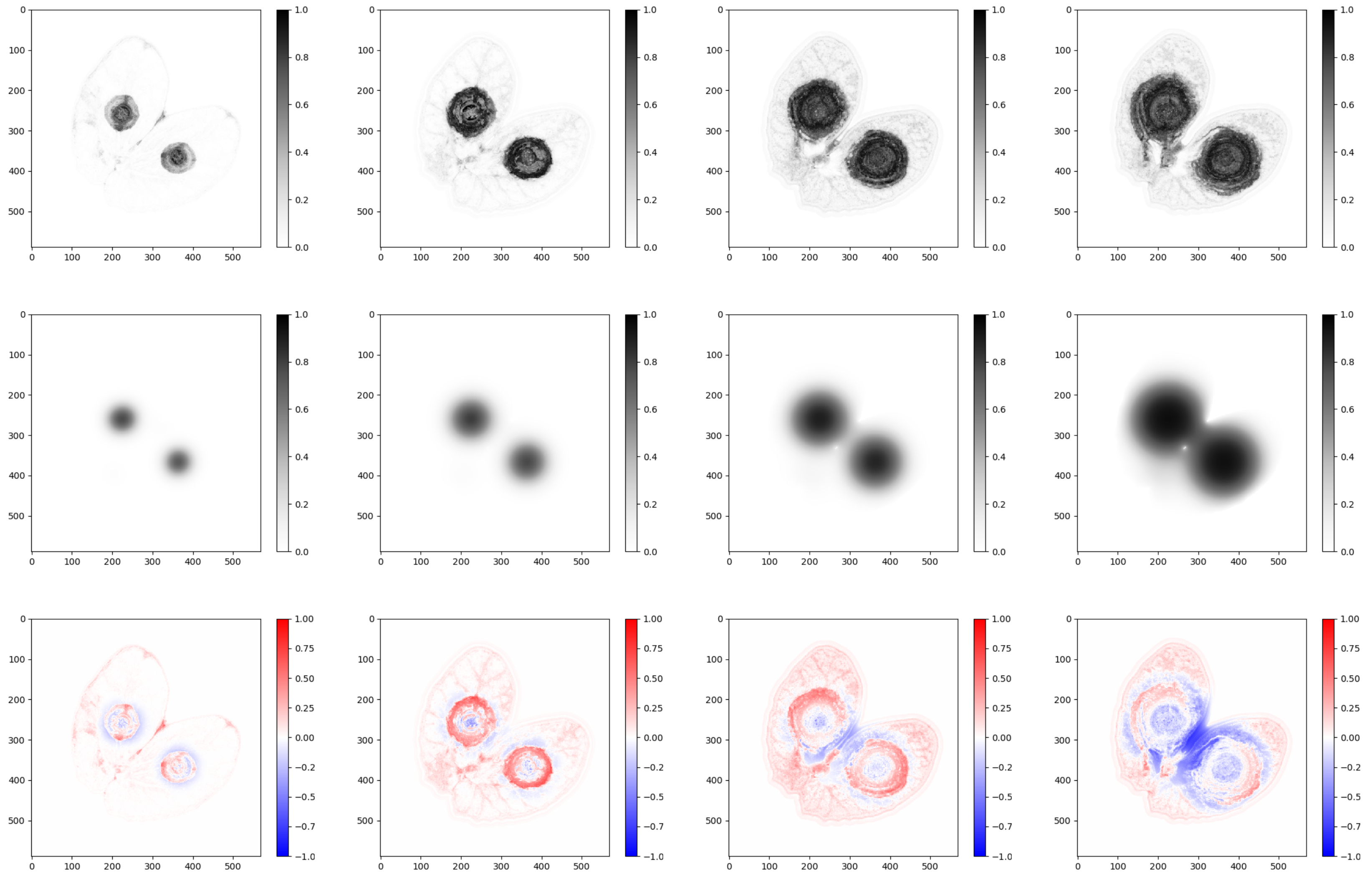

Figure F: Solara N°6

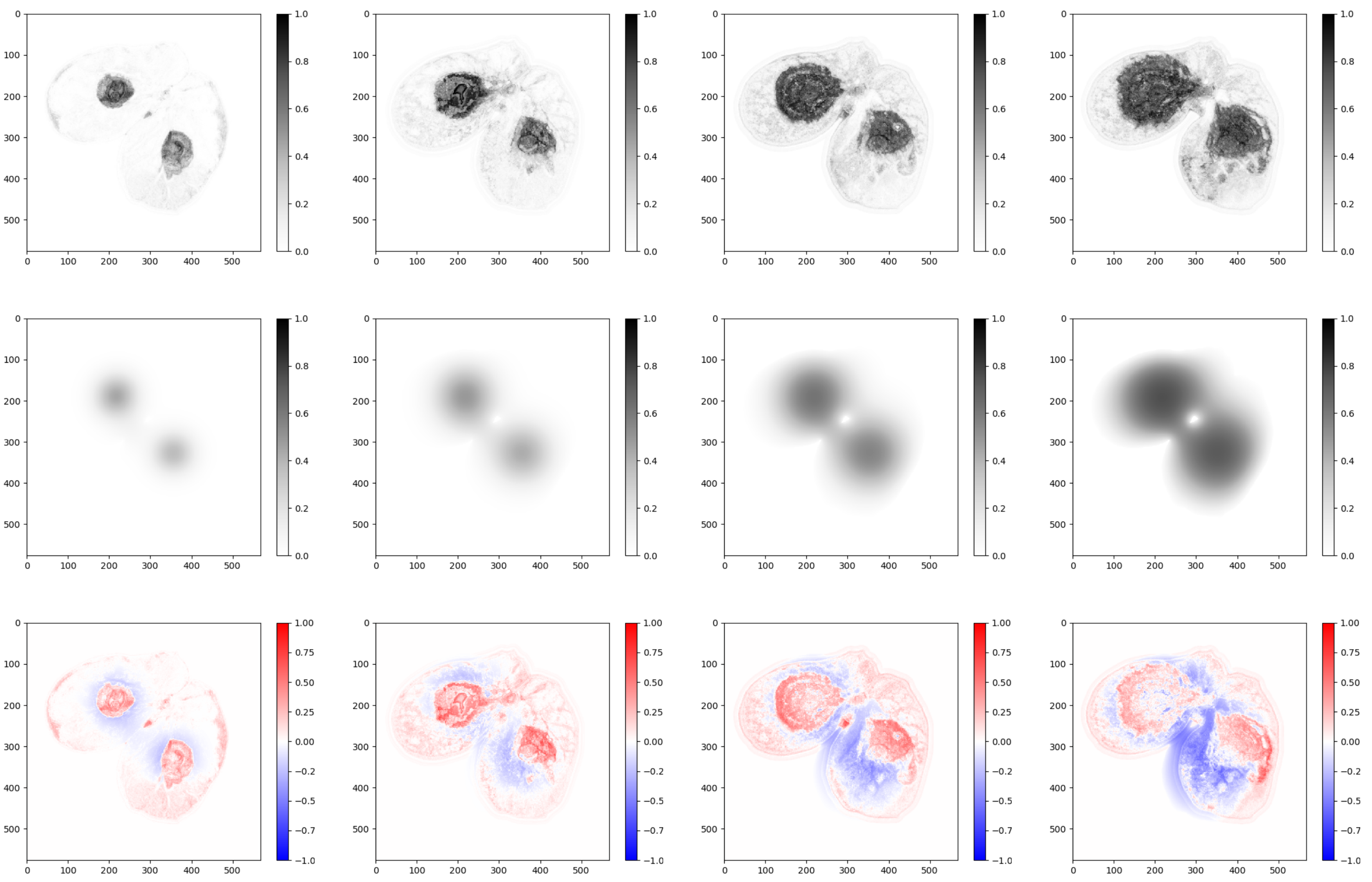

Figure G: Solara N°7

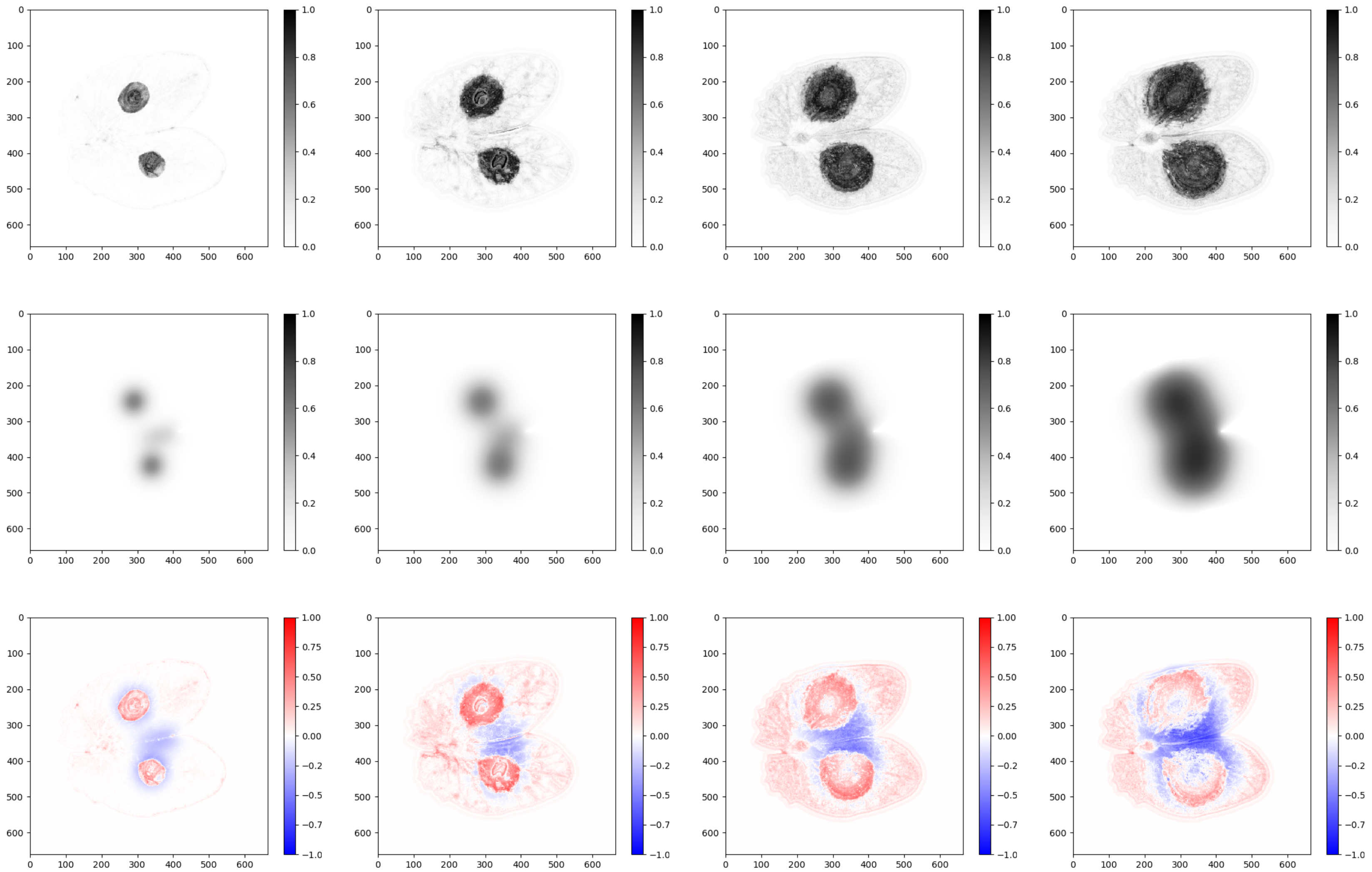


Figure H: Solara N°8

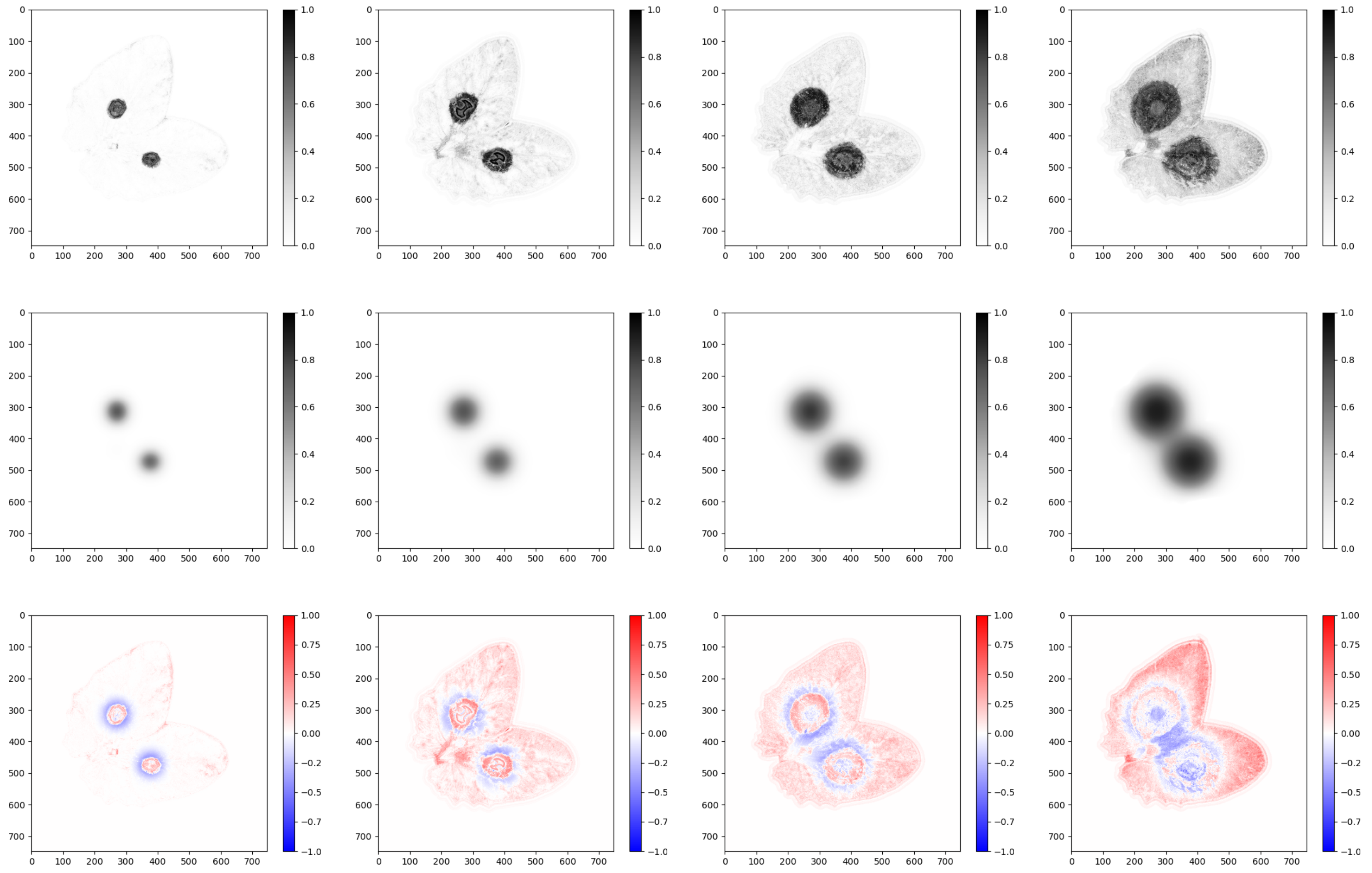

Figure I: Solara N°9

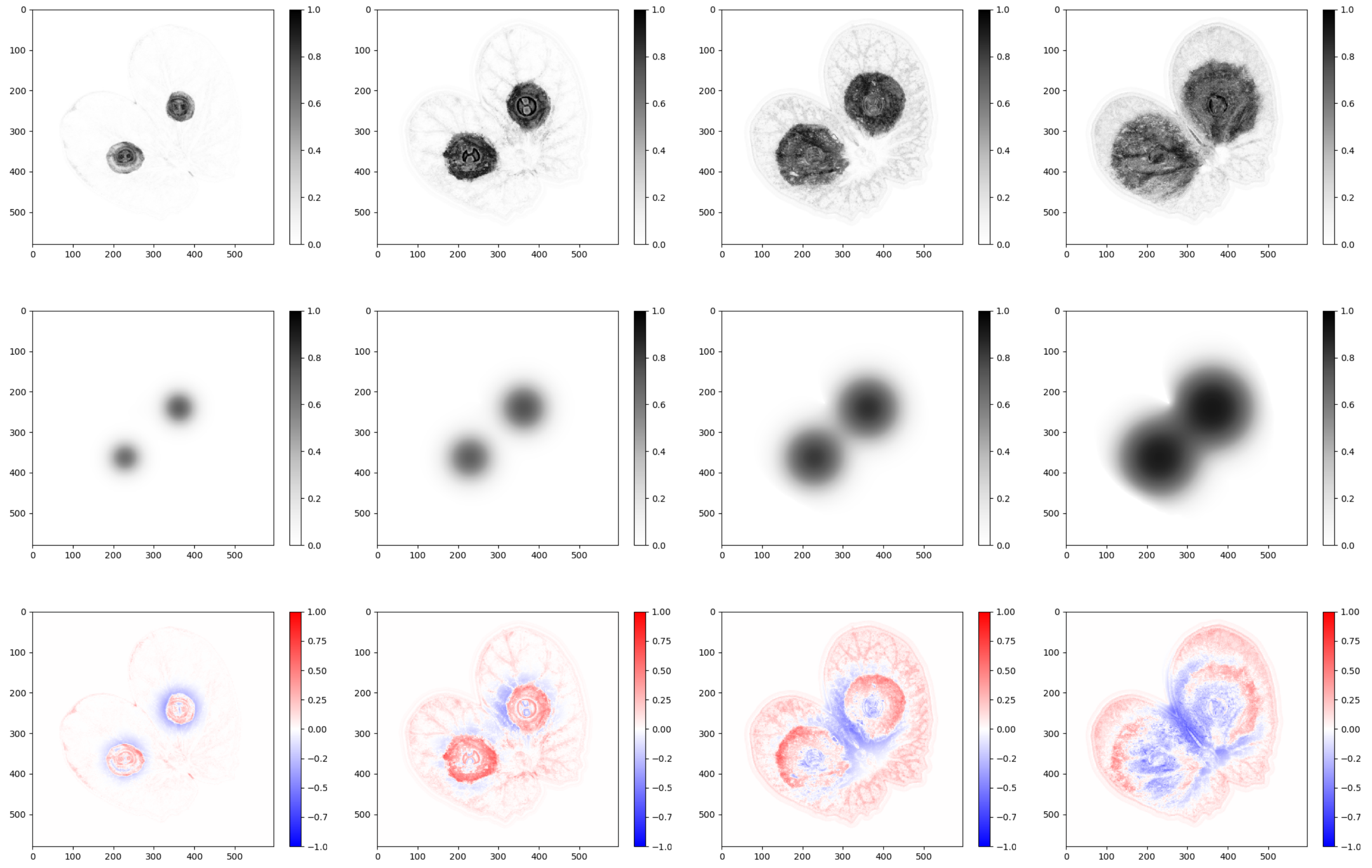

Figure J: Solara N°10

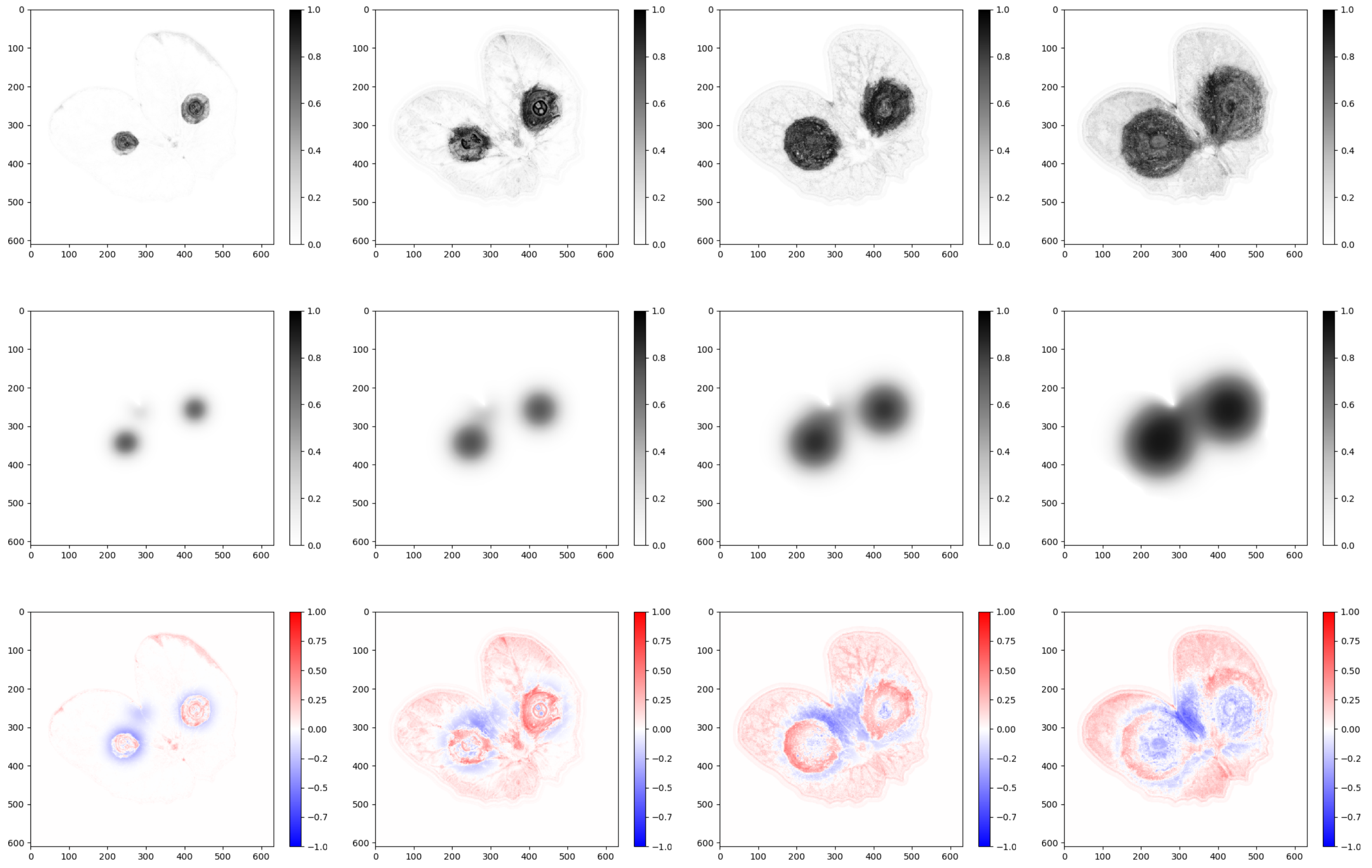


Figure K: Solara N°11

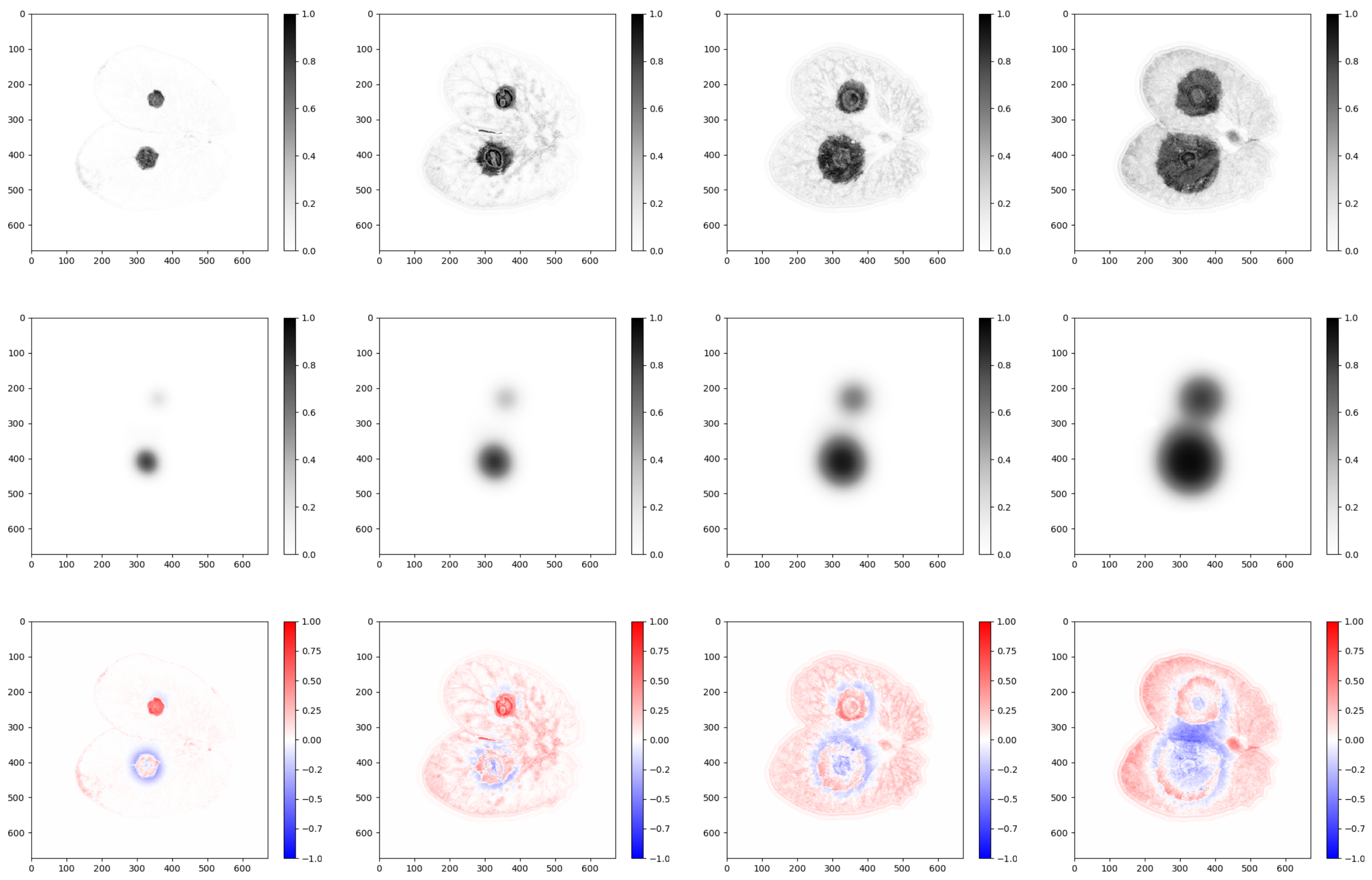

Figure L: Solara N°12

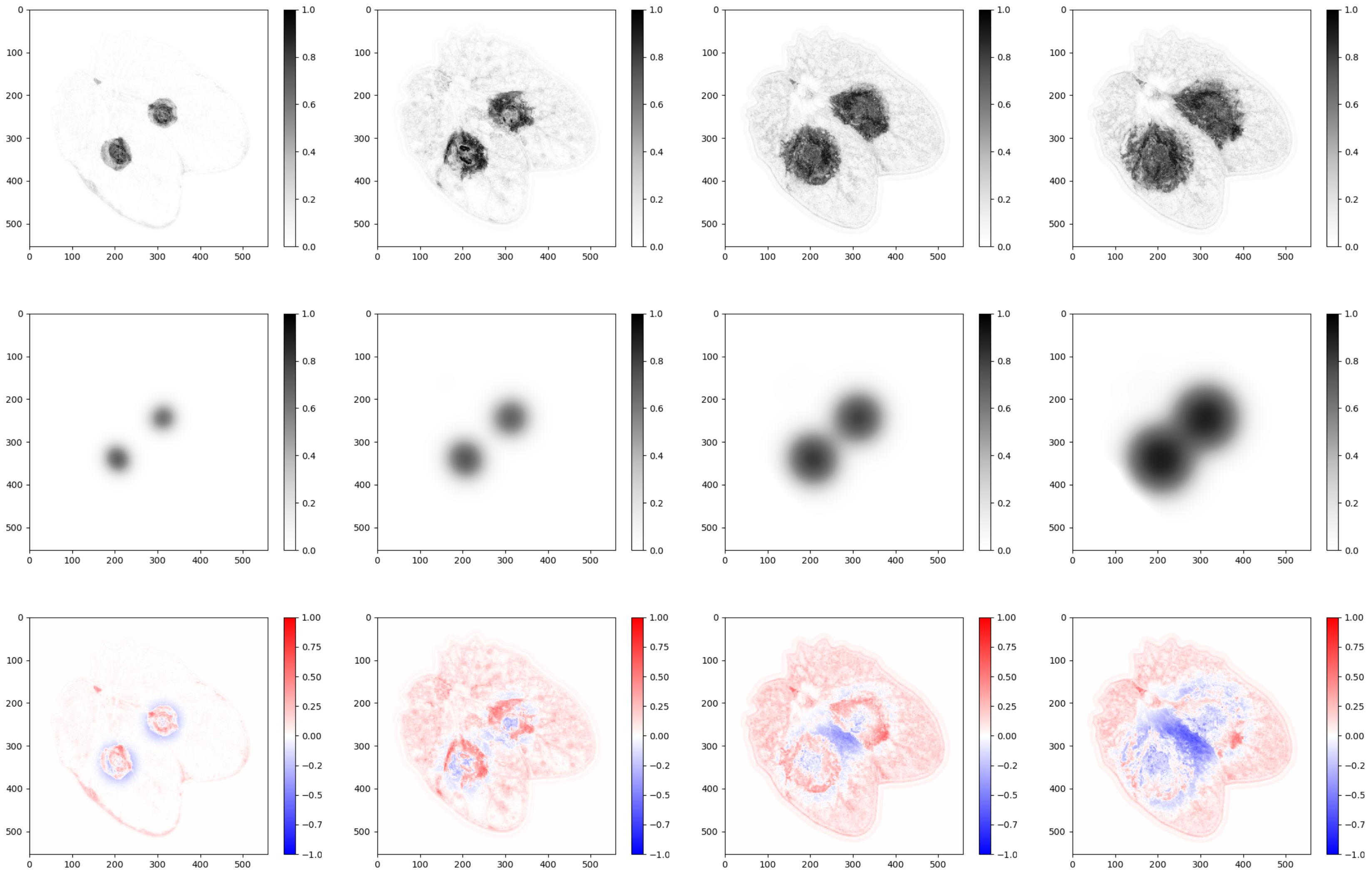

Figure M: Solara N°13

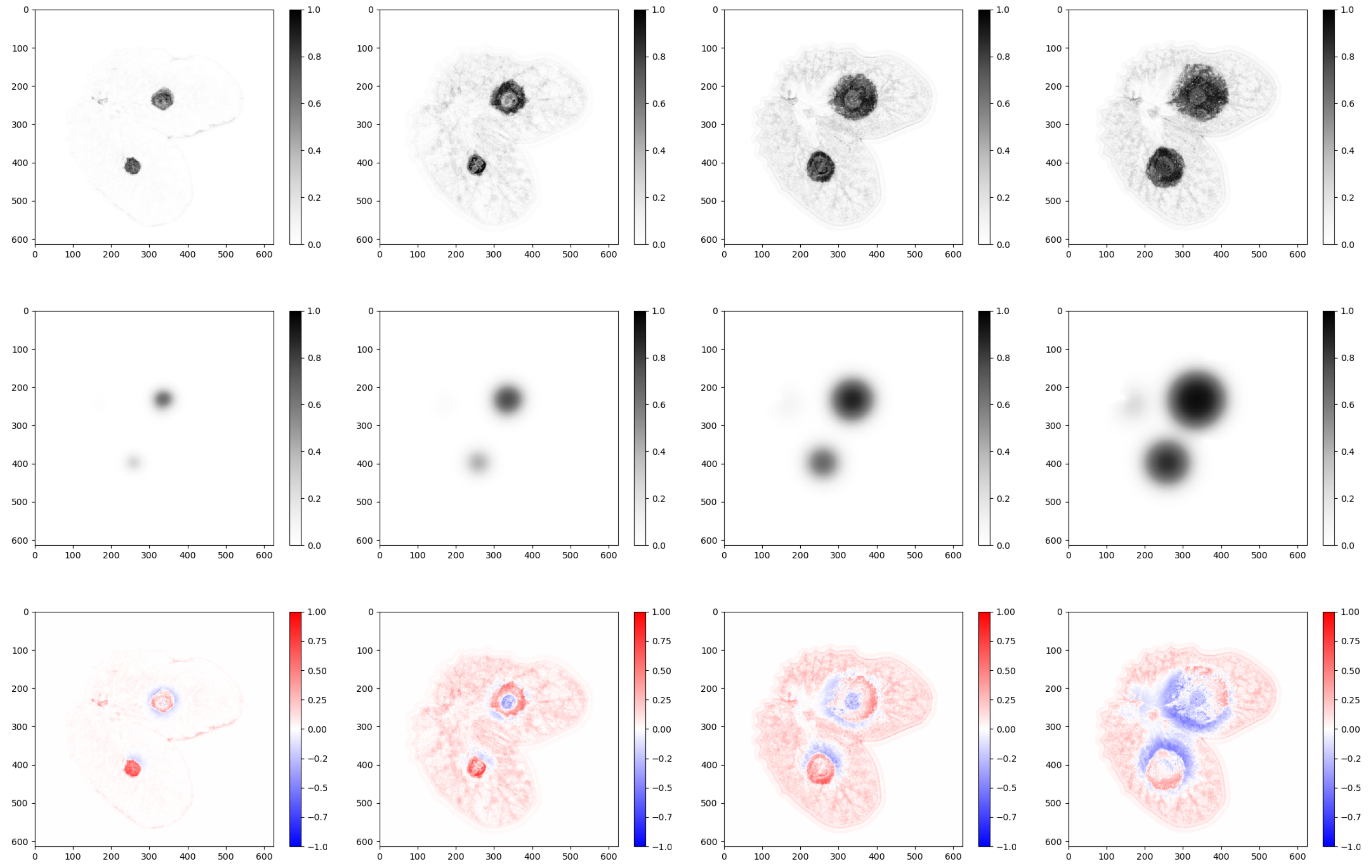

Figure N: Solara N°14

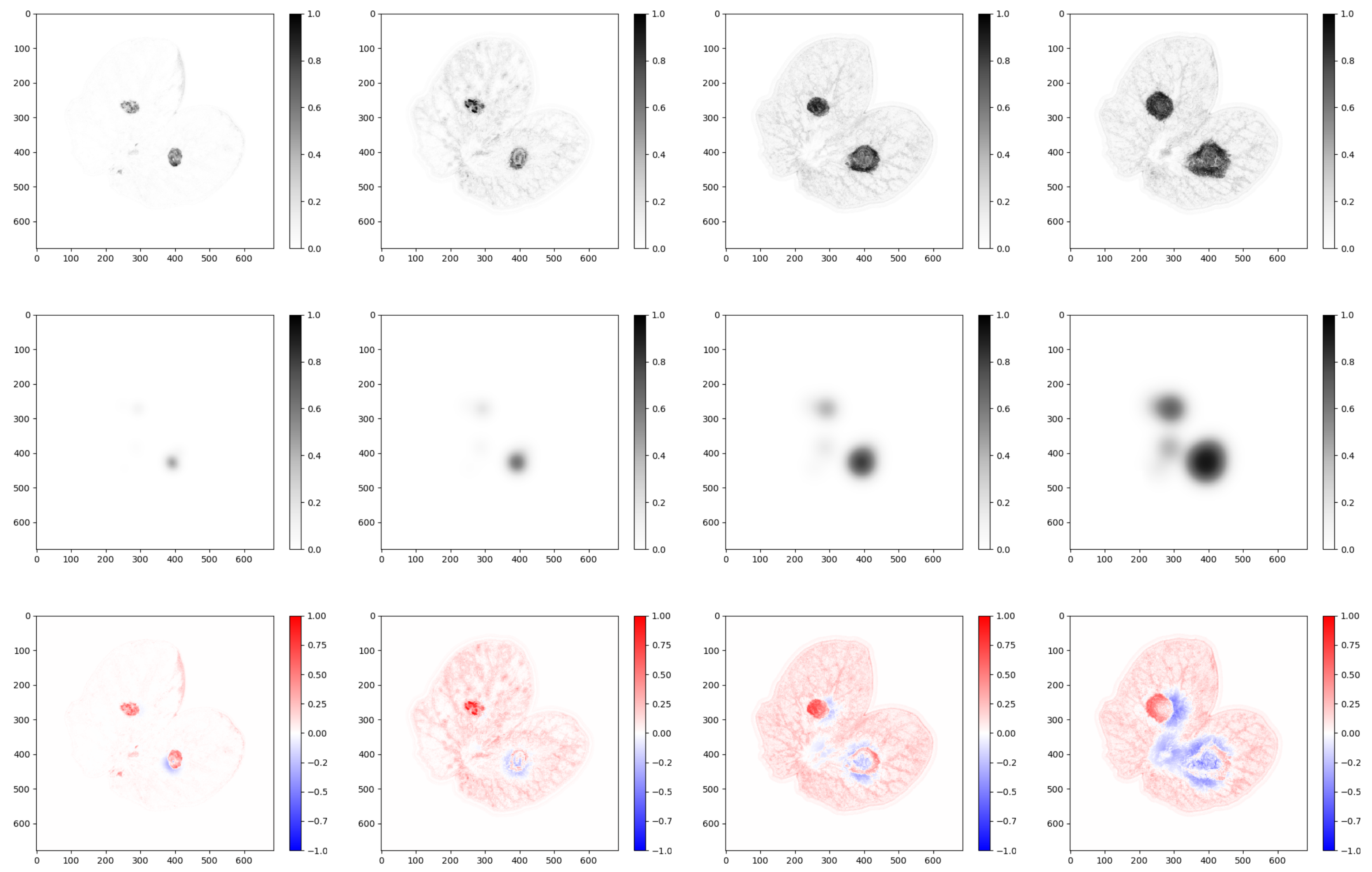


Figure O: Solara N°15

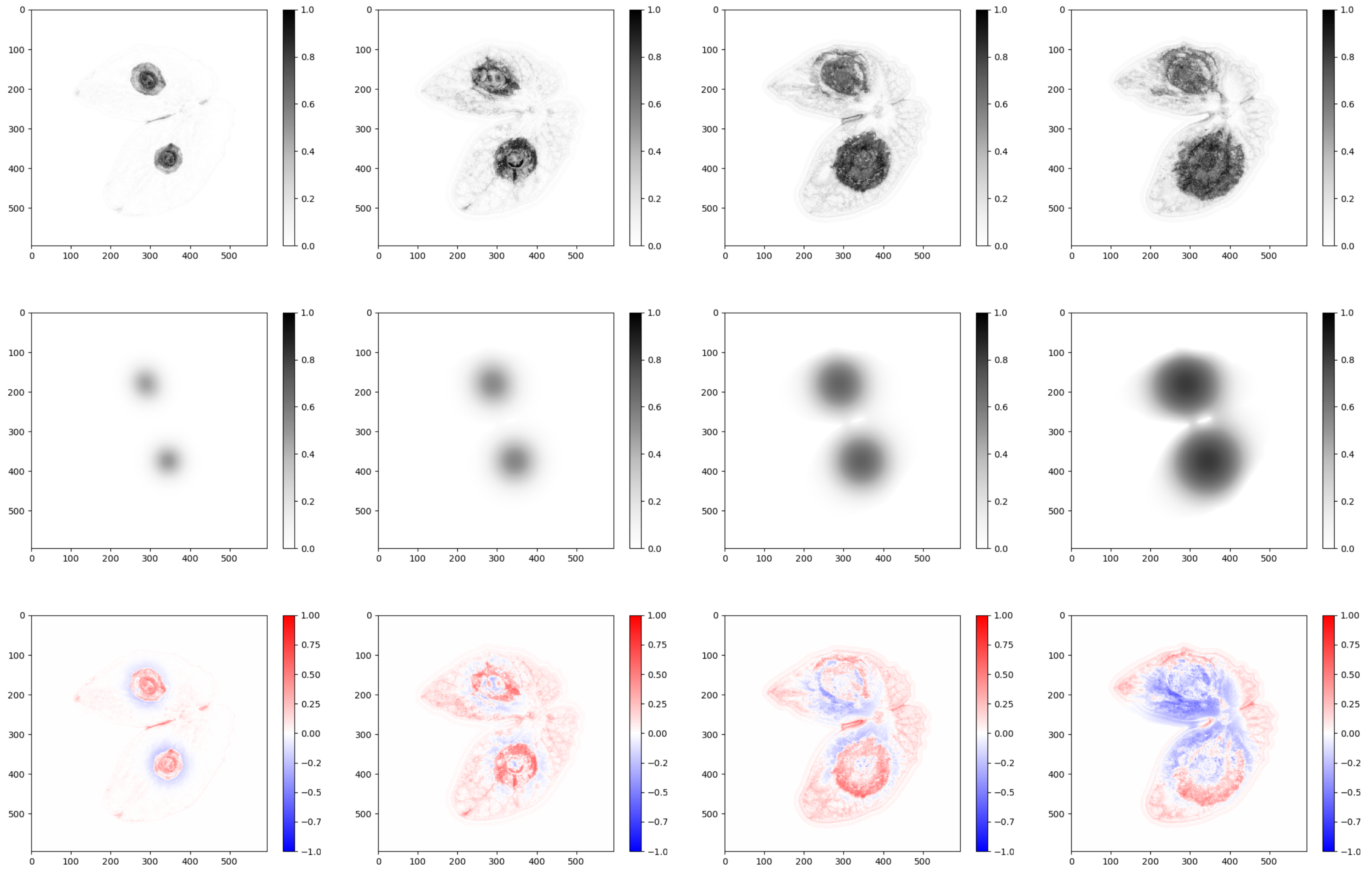

Figure P: Solara N°16

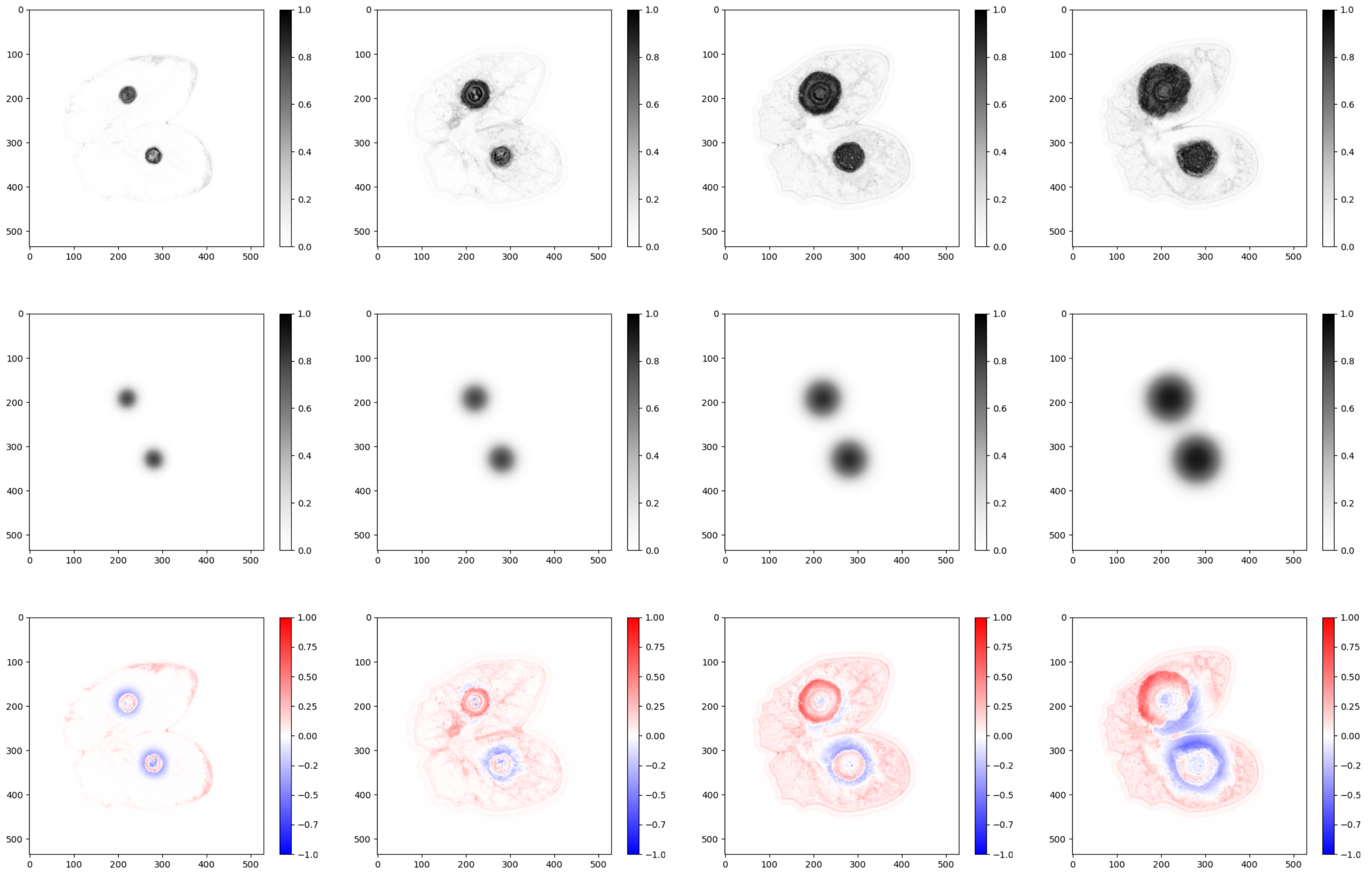

Figure Q: James N°17

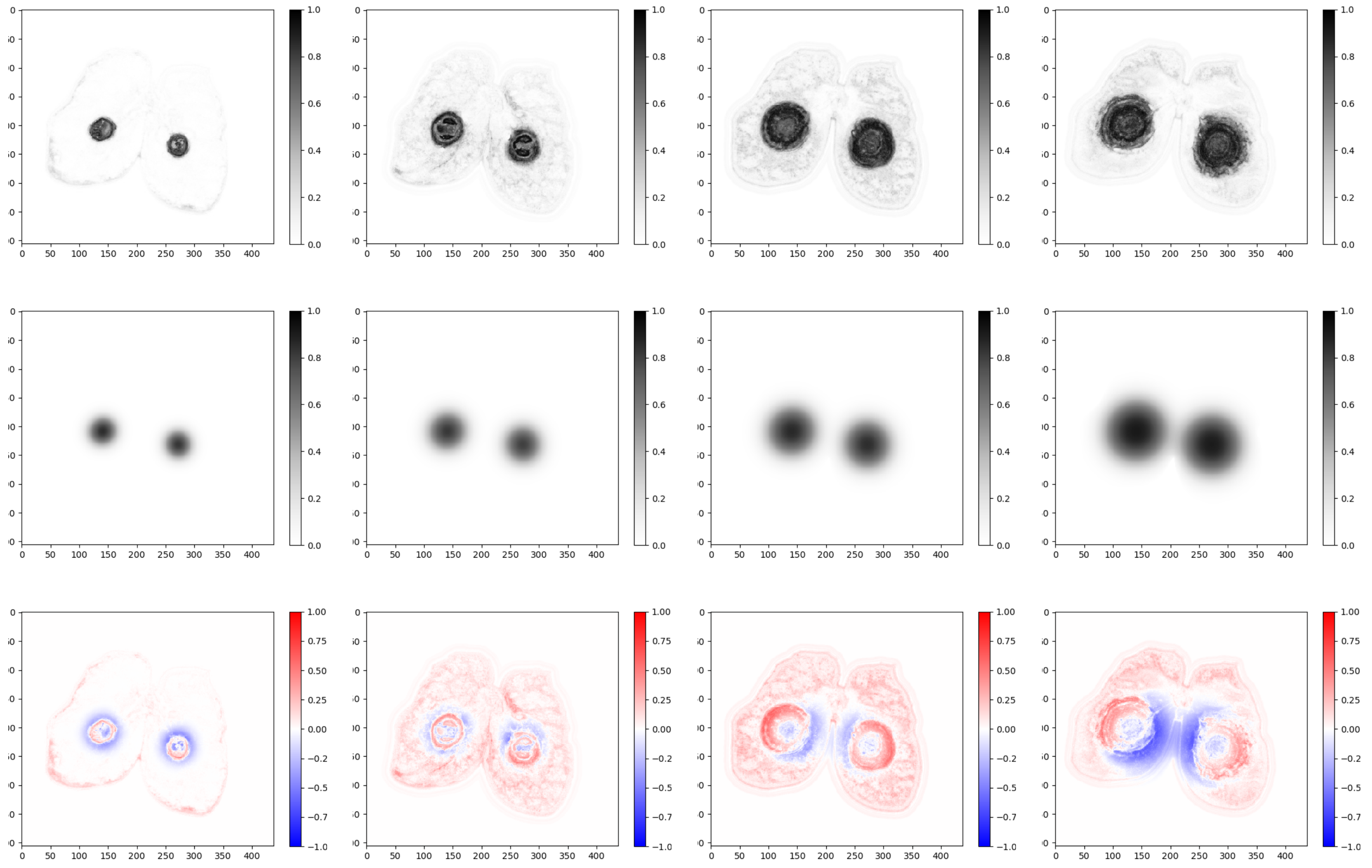

Figure R: James N°18

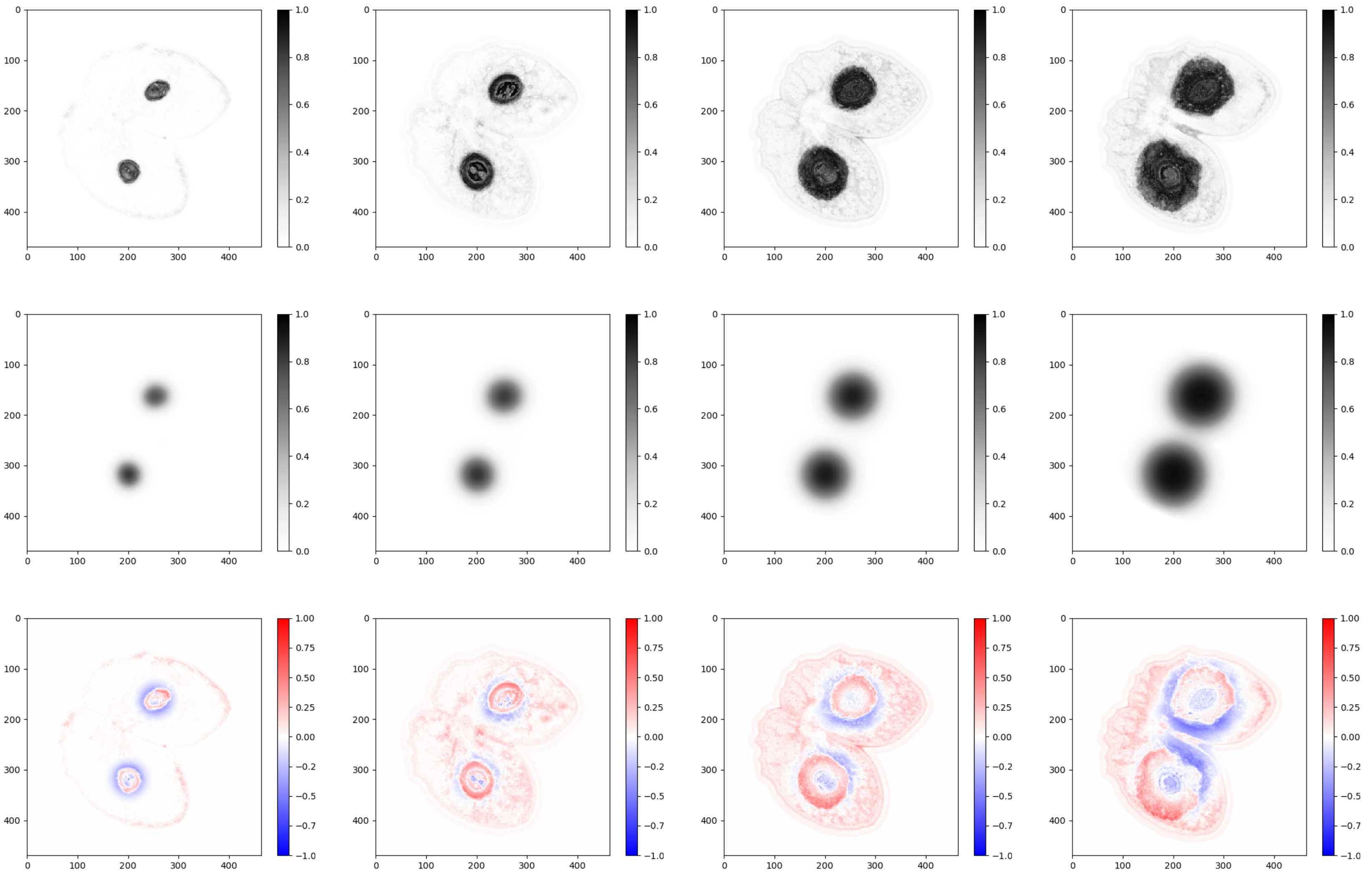

Figure S: James N°19

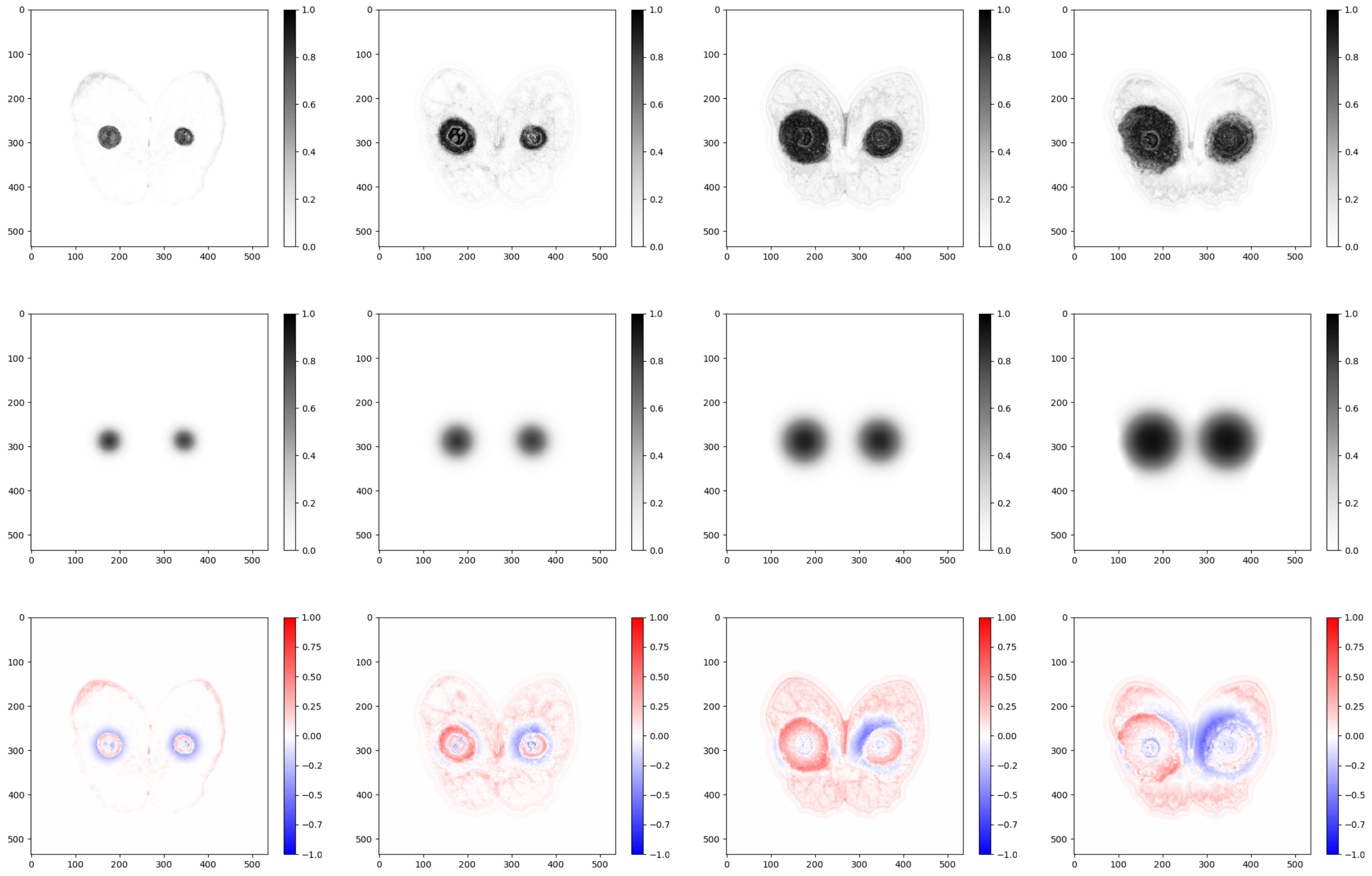

Figure T: James N°20

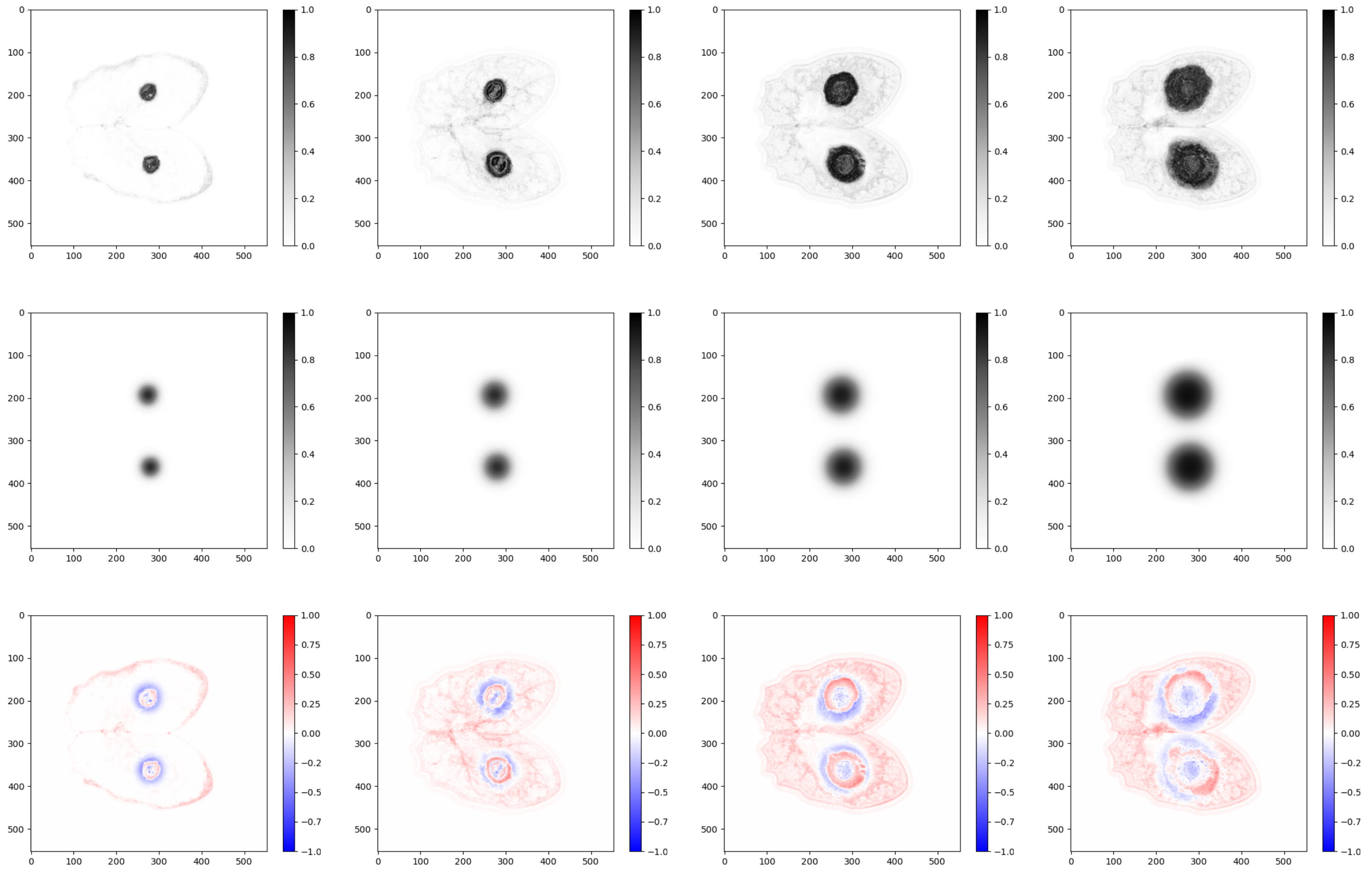

Figure U: James N°21

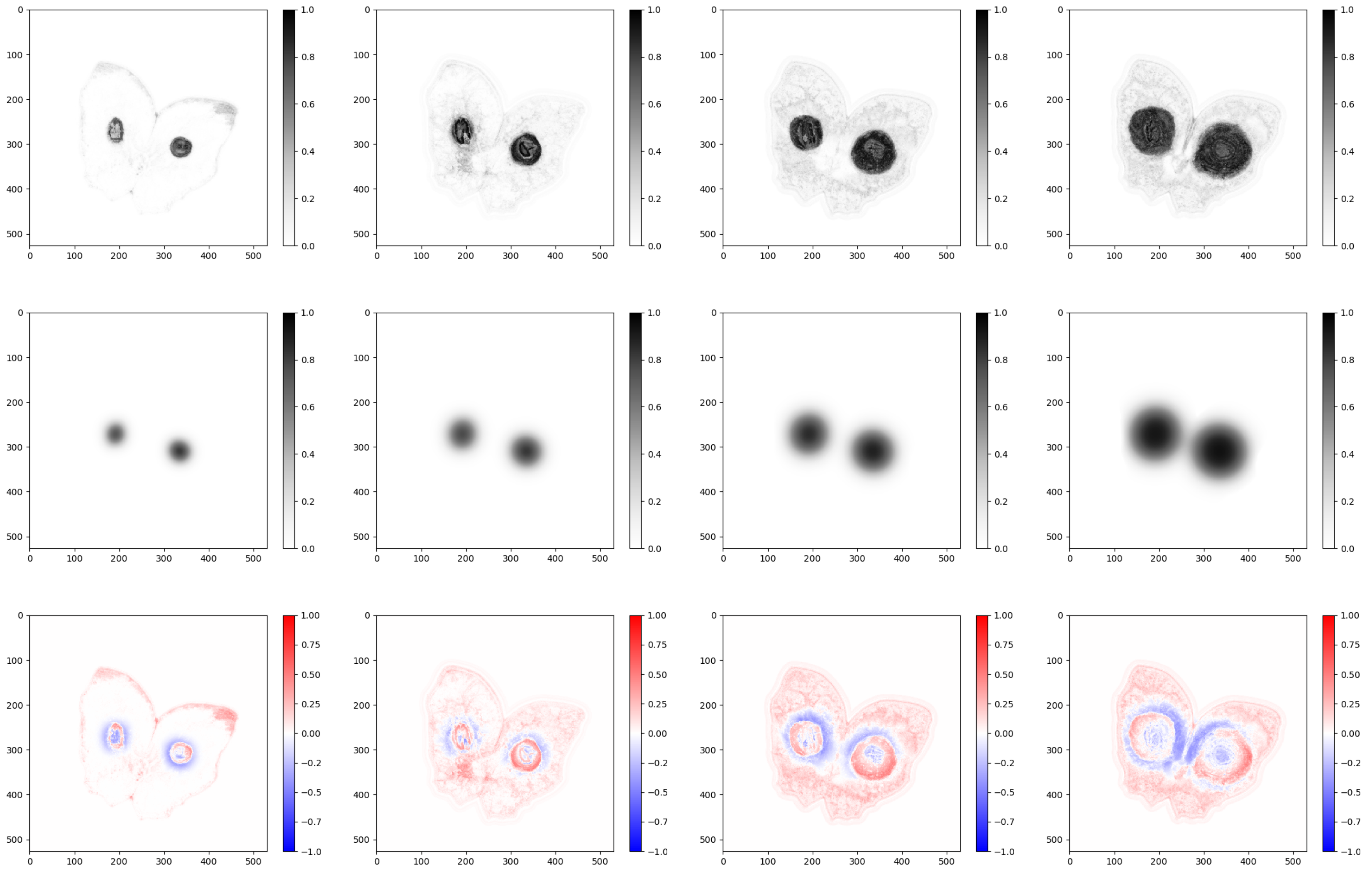

Figure V: James N°22

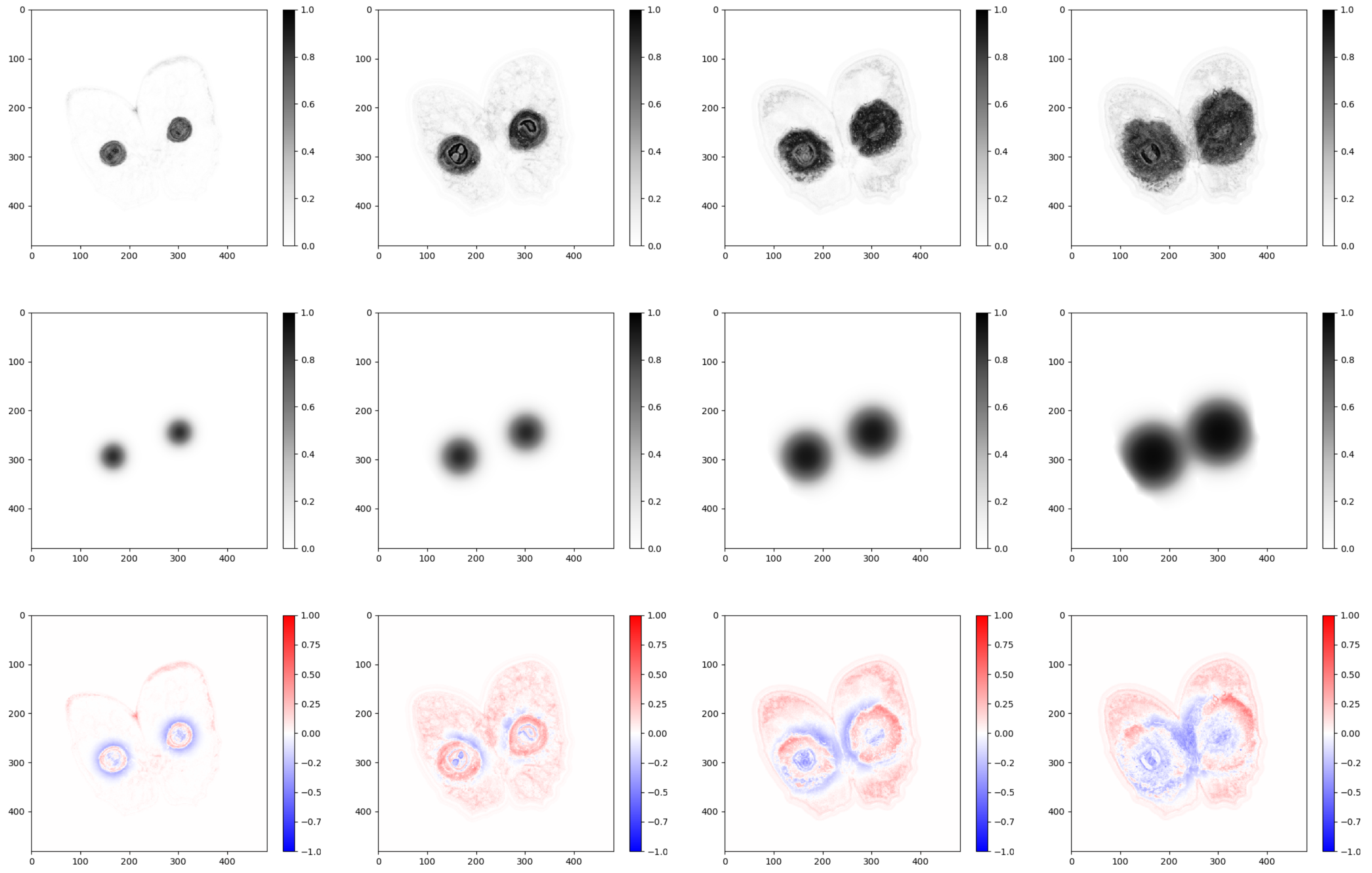

Figure W: James N°23

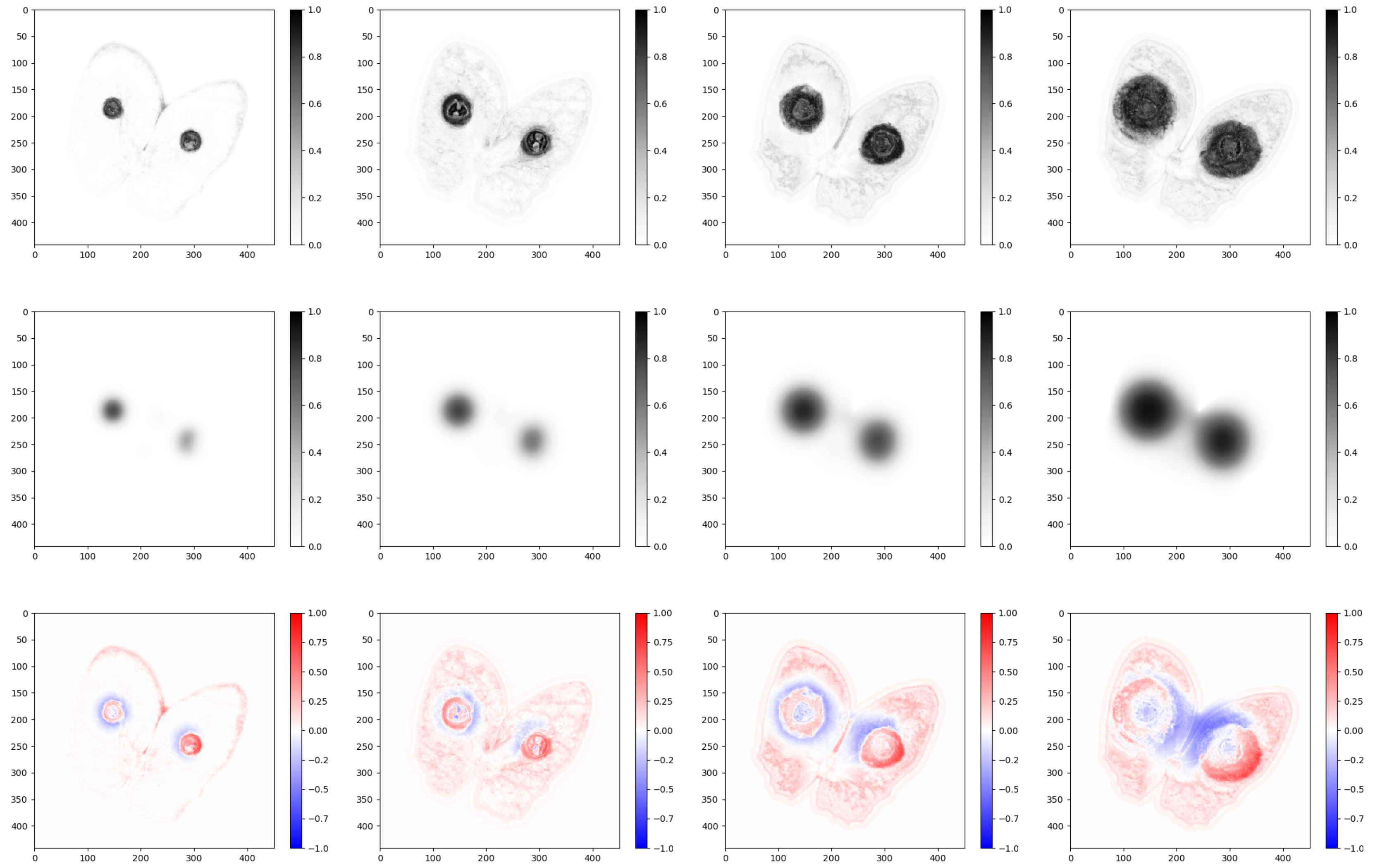


Figure X: James N°24

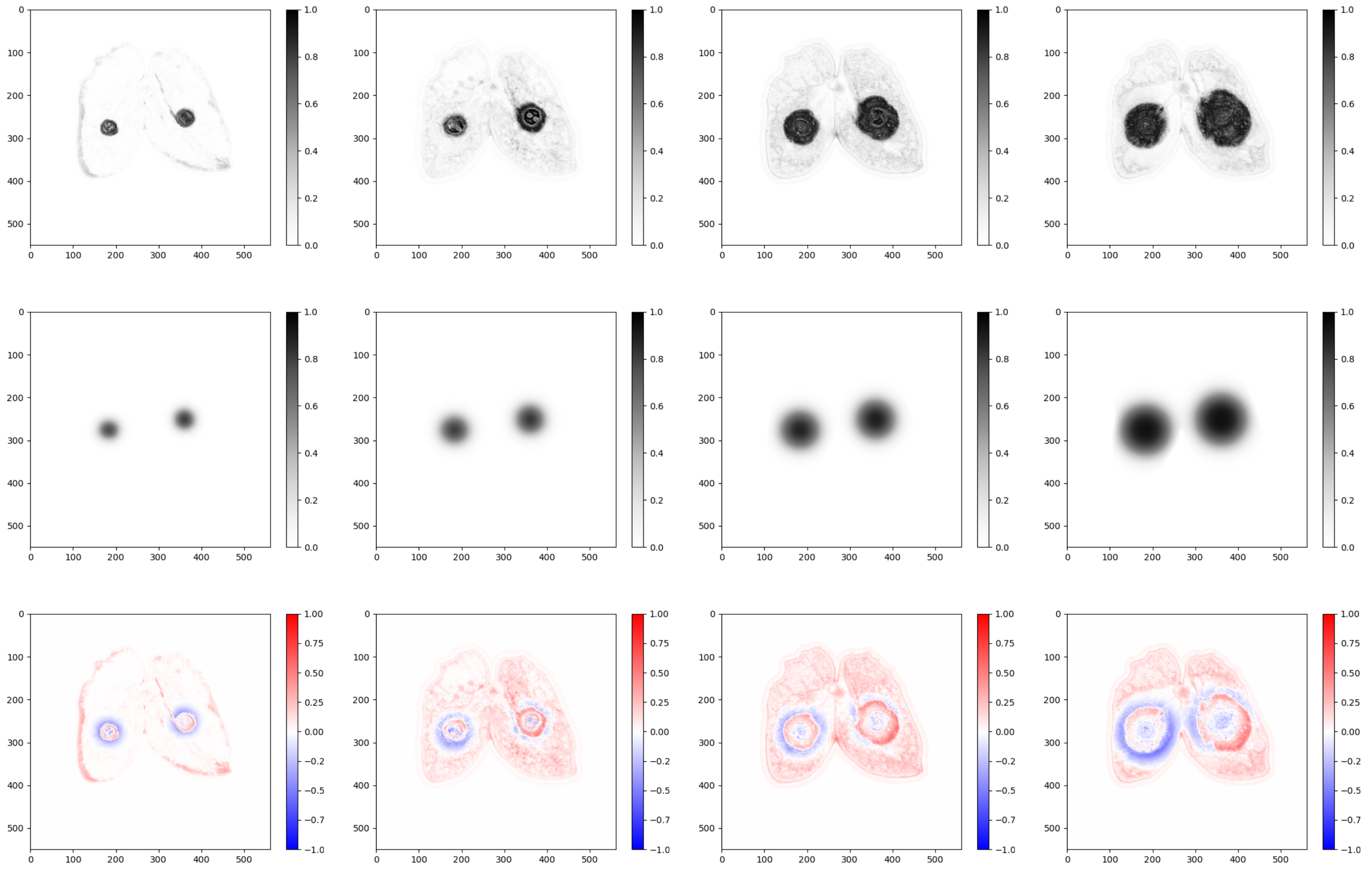

Figure Y: James N°25

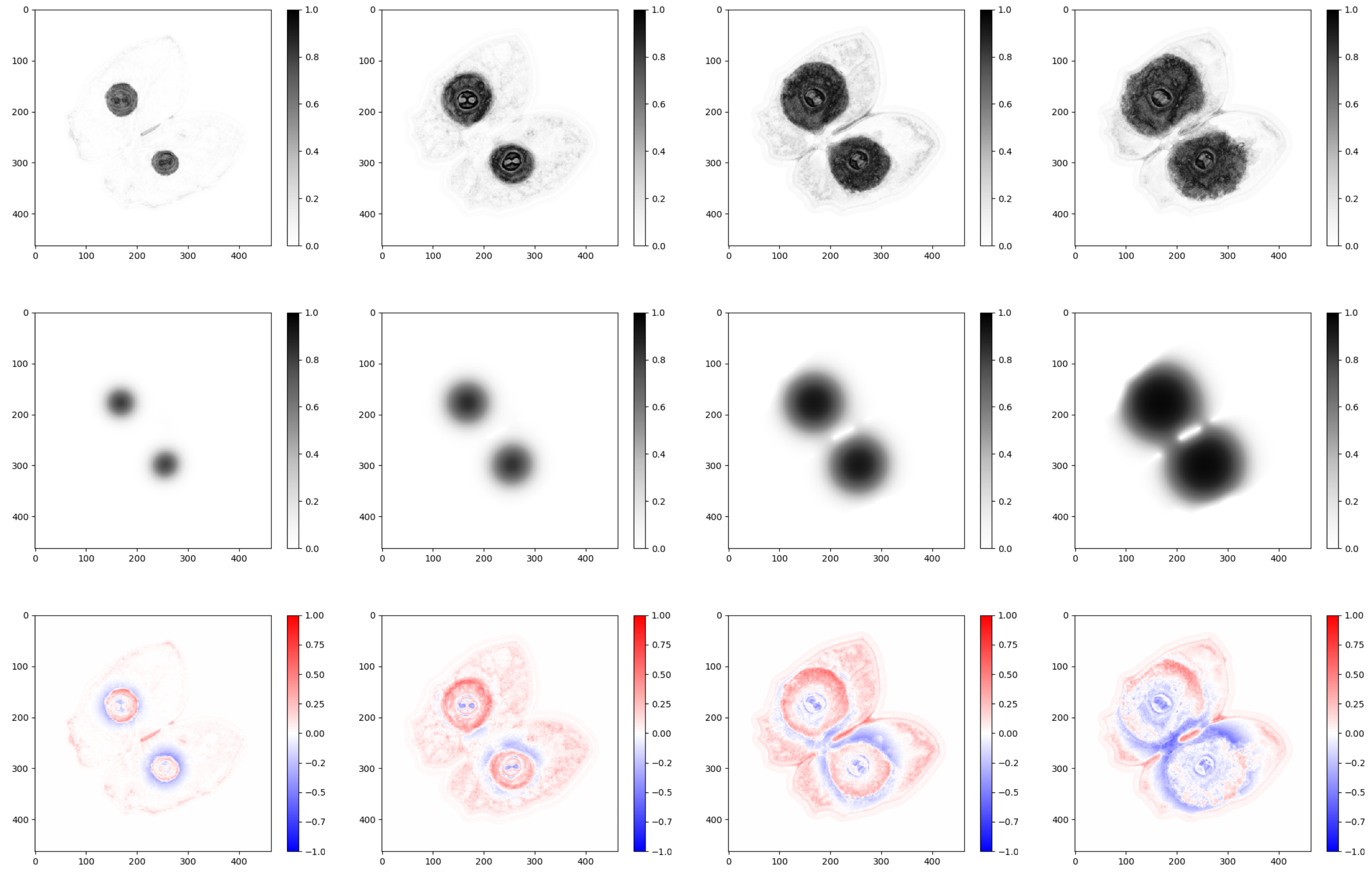


Figure Z: James N°26

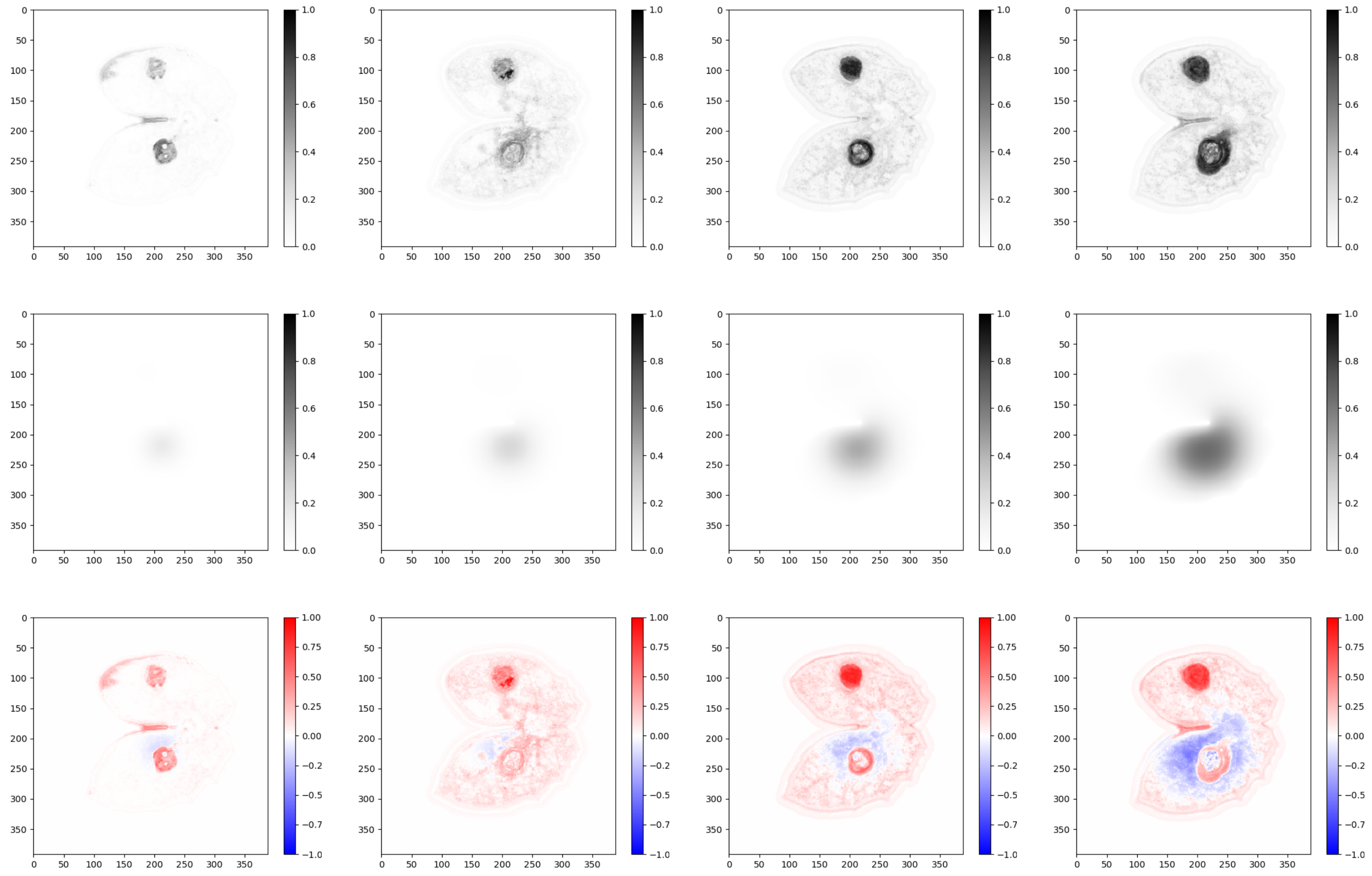

Figure AA: James N°27

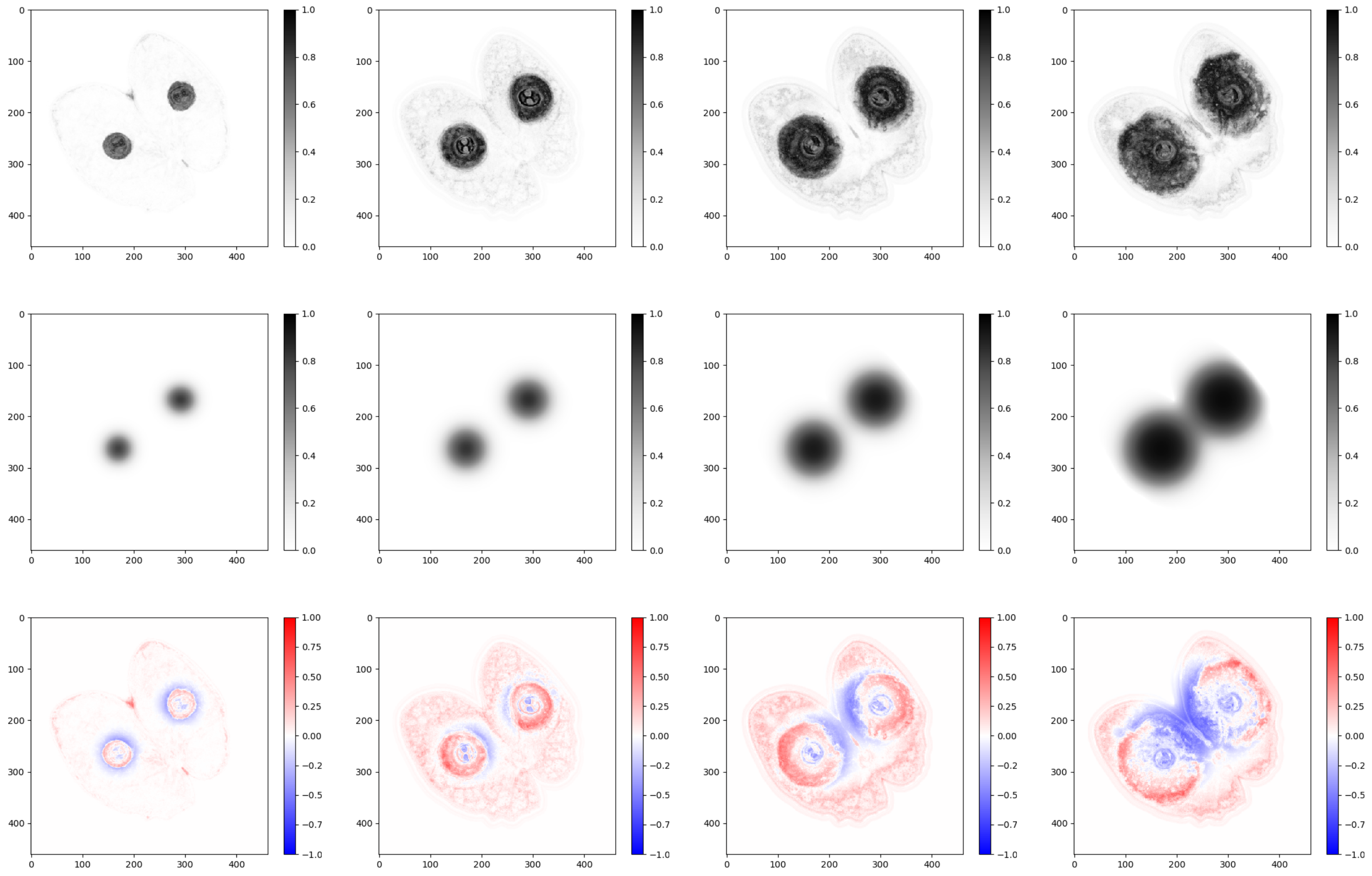

Figure AB: James N°28

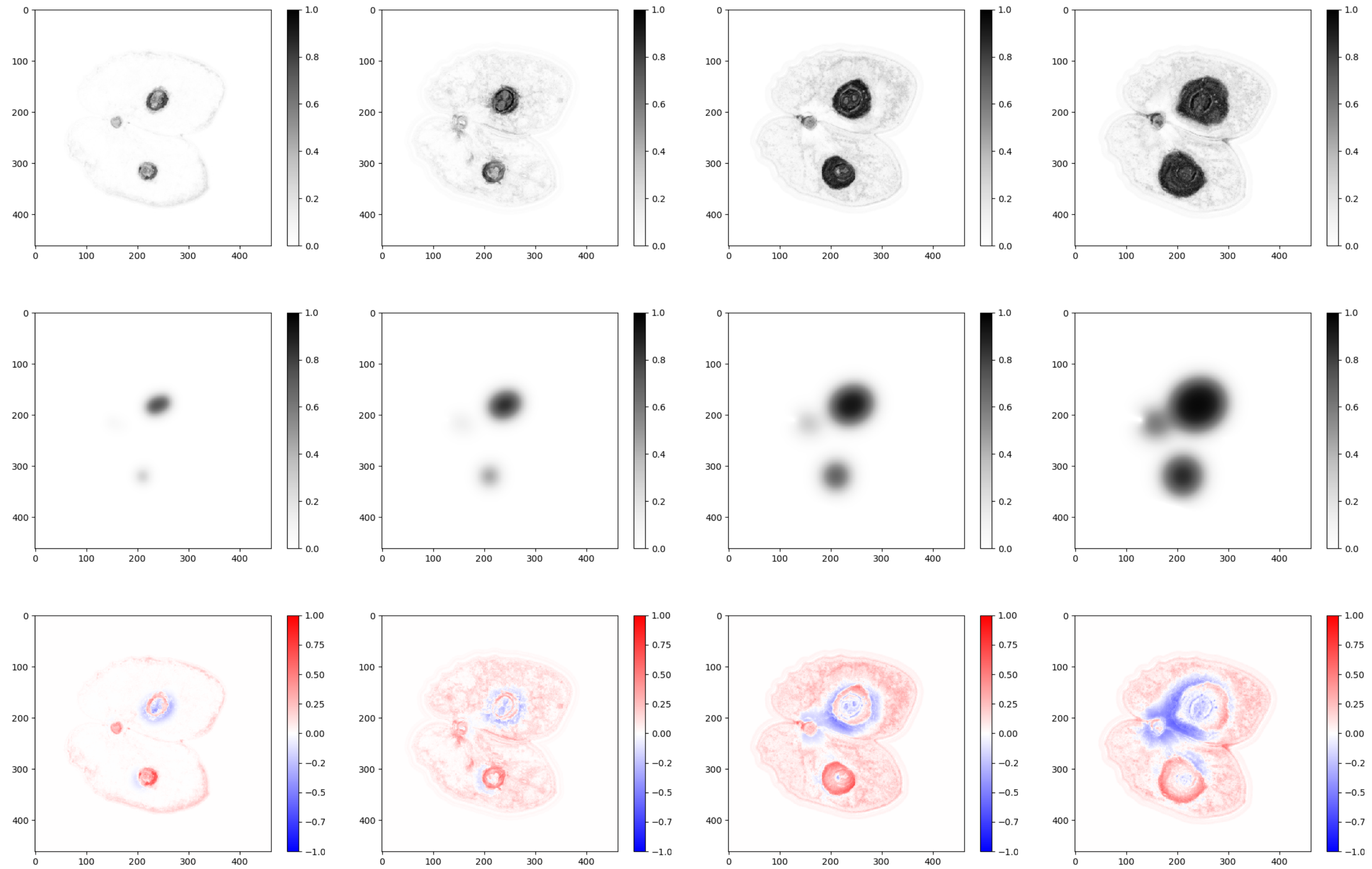

Figure AC: James N°29

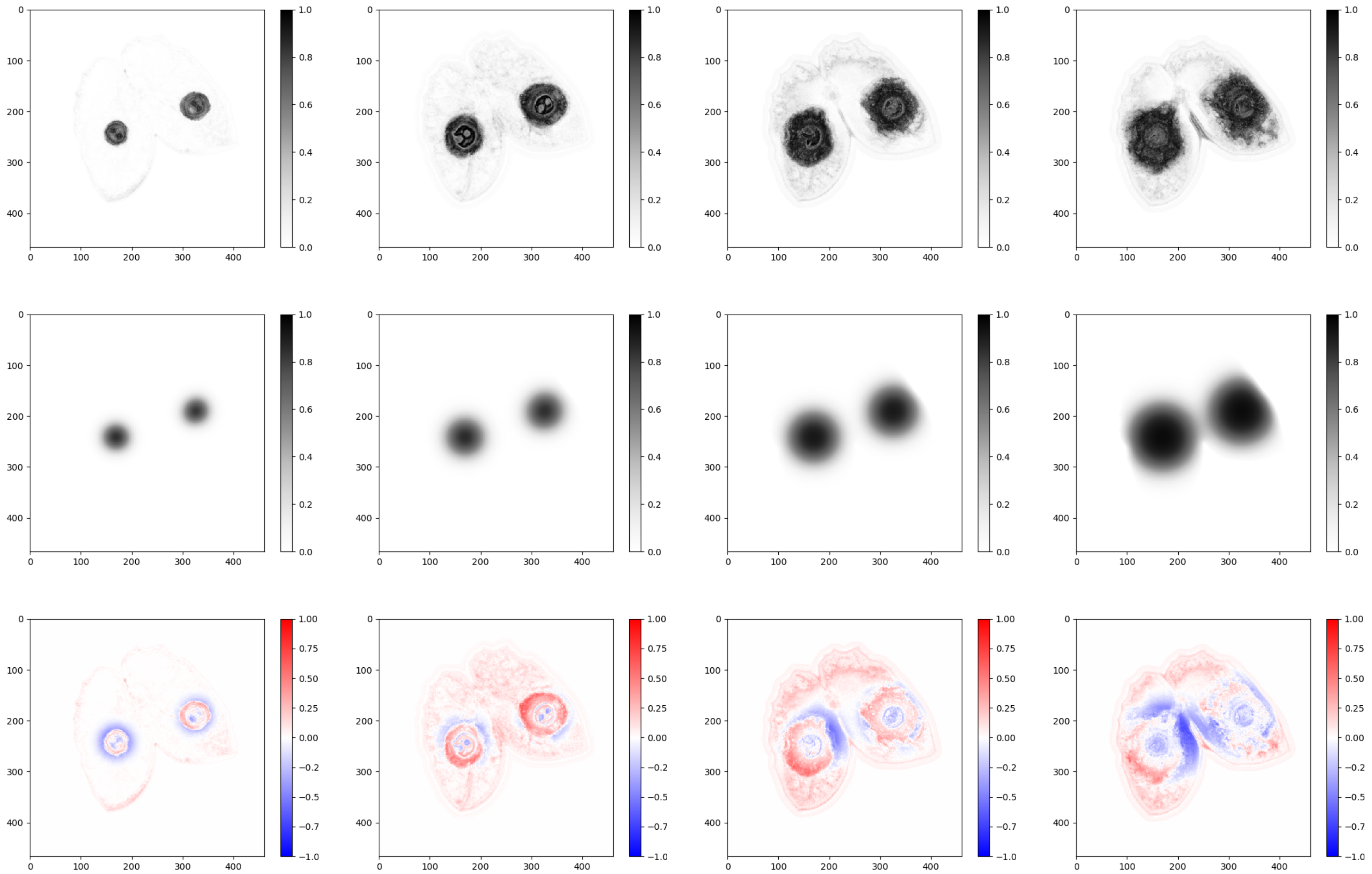

Figure AD: James N°30

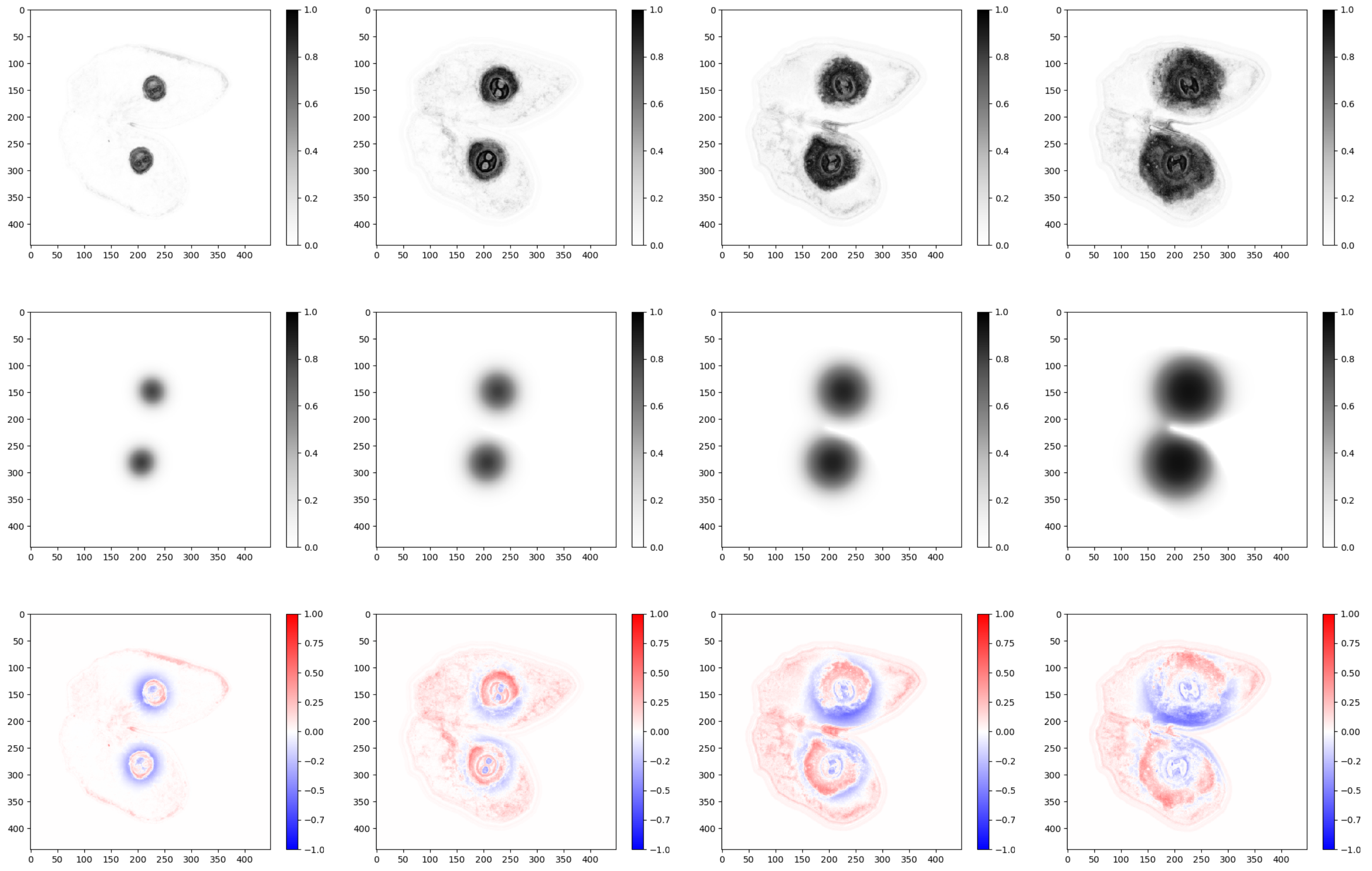

Figure AE: James N°31

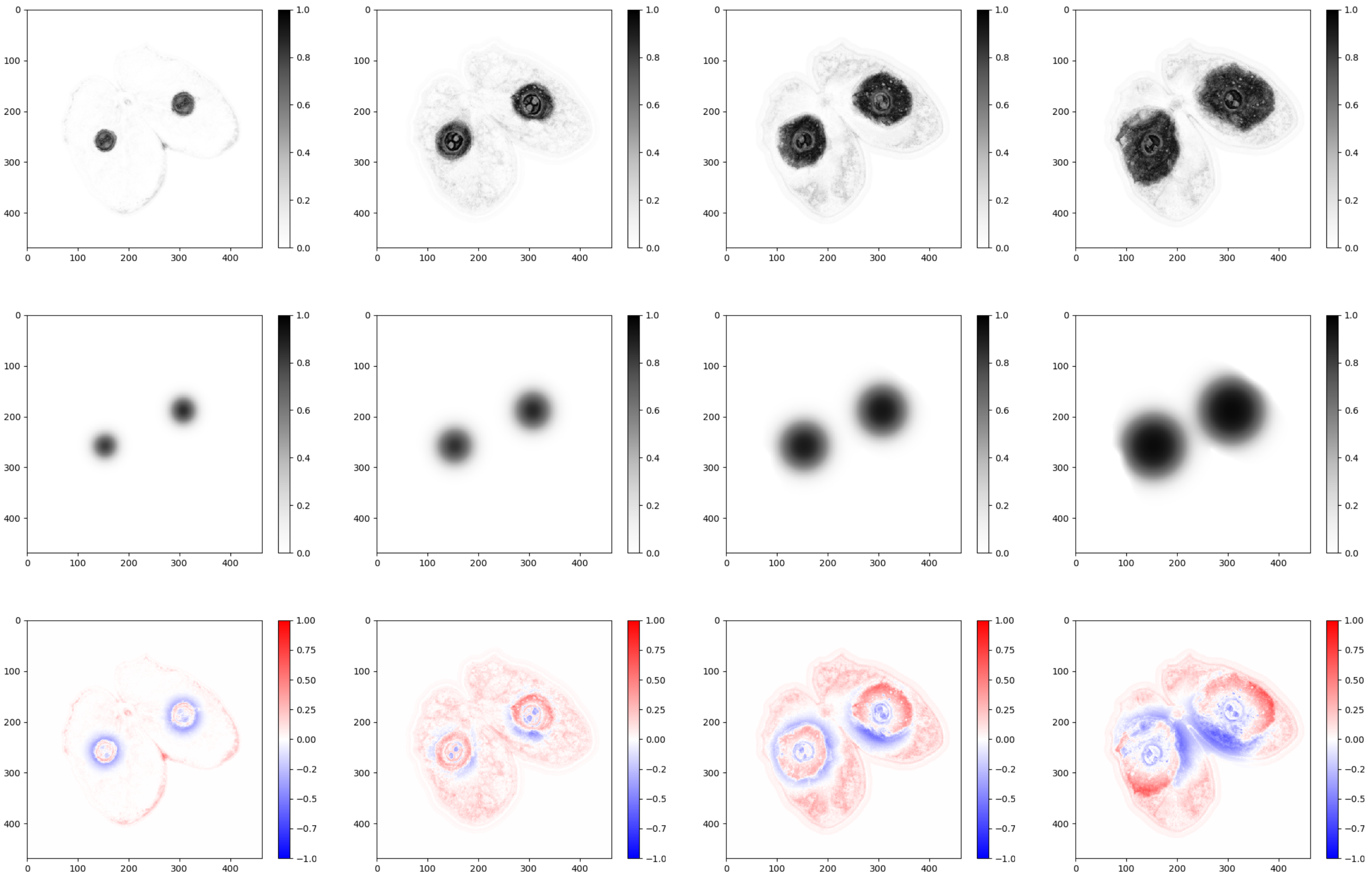

Figure AF: James N°32

#### S3.2 Visualization of raw residuals

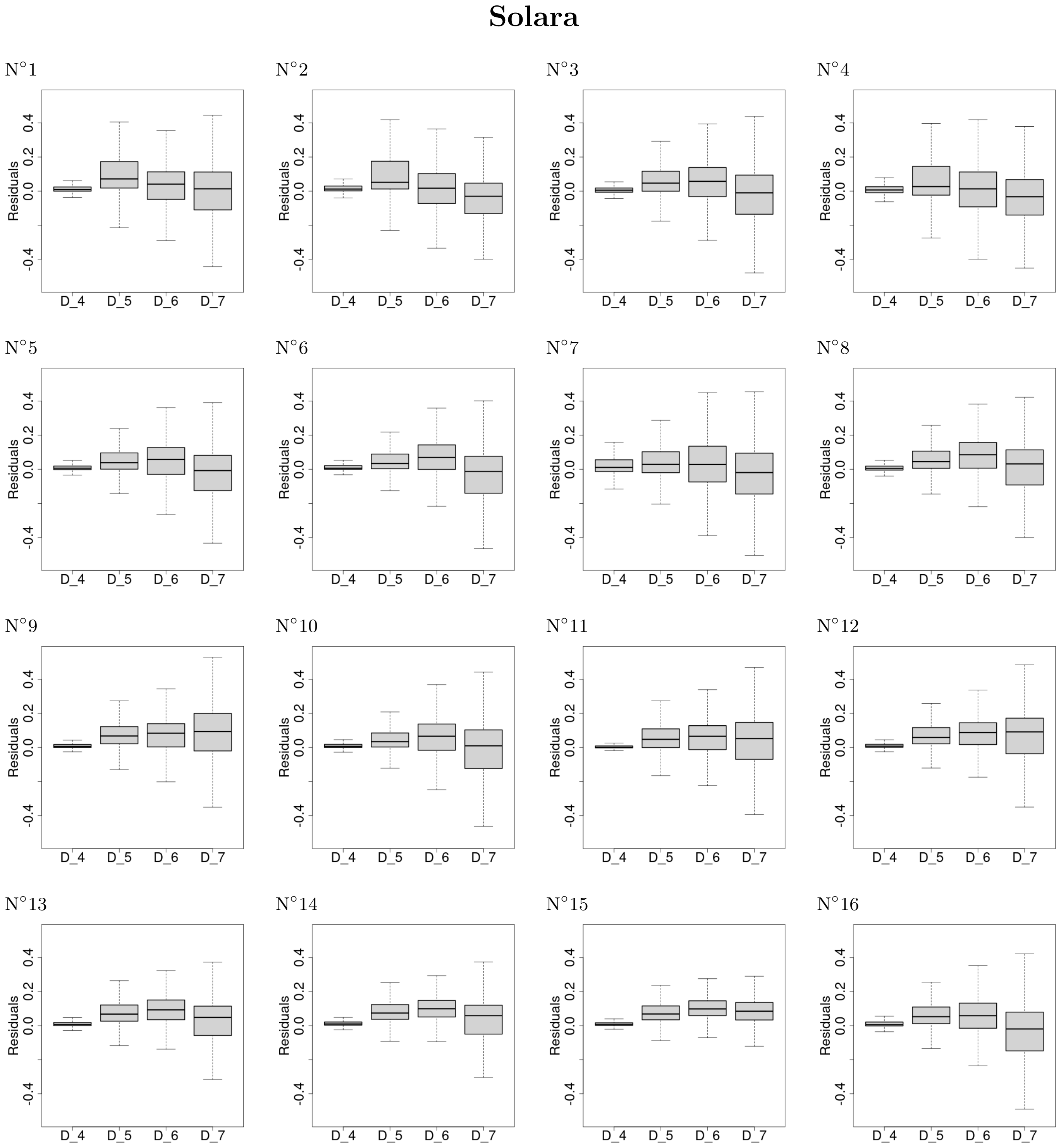


Figure AG: Distribution of raw residuals against dates after inoculation for each Solara stipule pairs.

### James

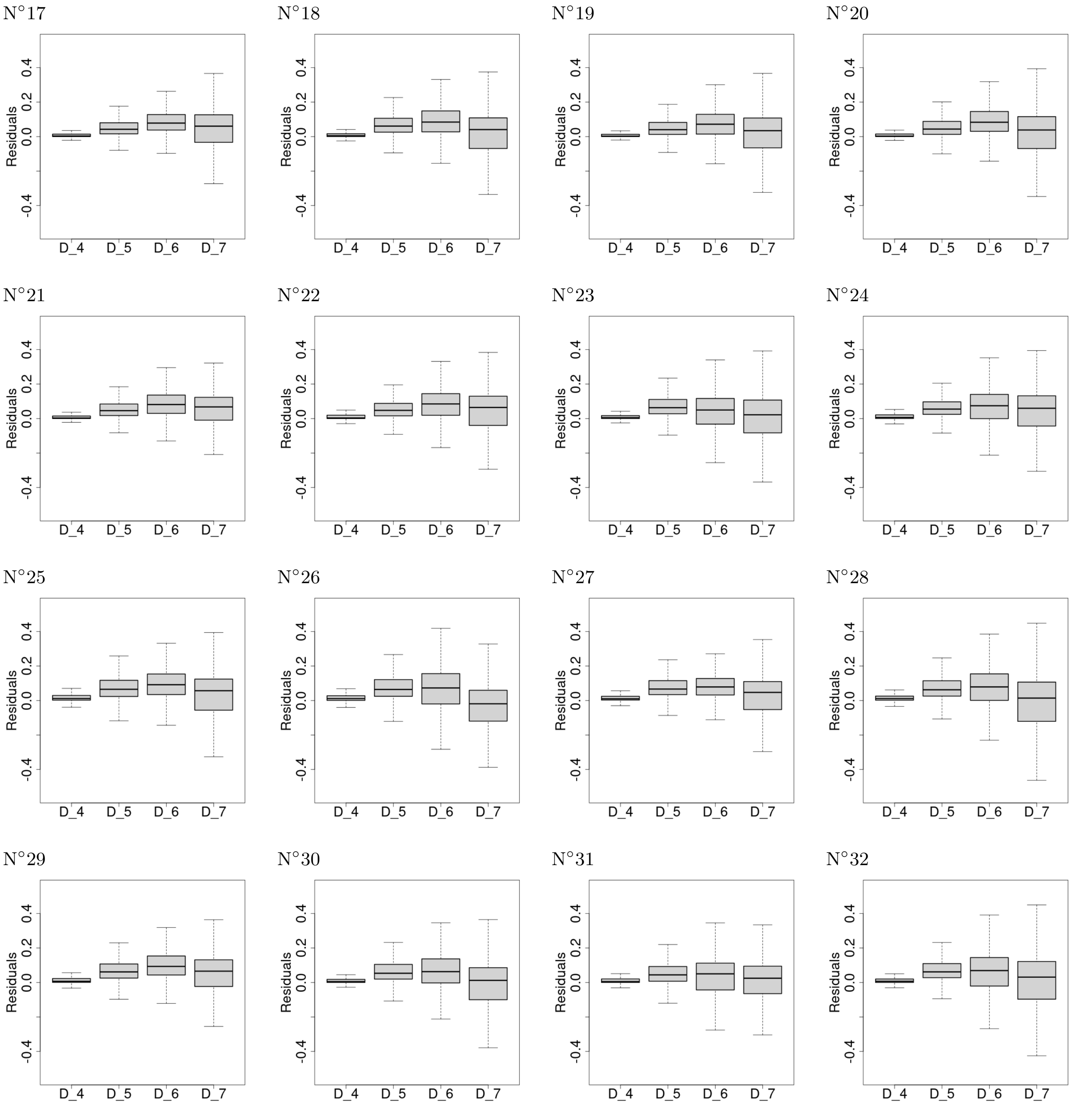


Figure AH: Distribution of raw residuals against dates after inoculation for each James stipule pairs.

#### S3.3 Model prediction against data (pixel versus pixel)

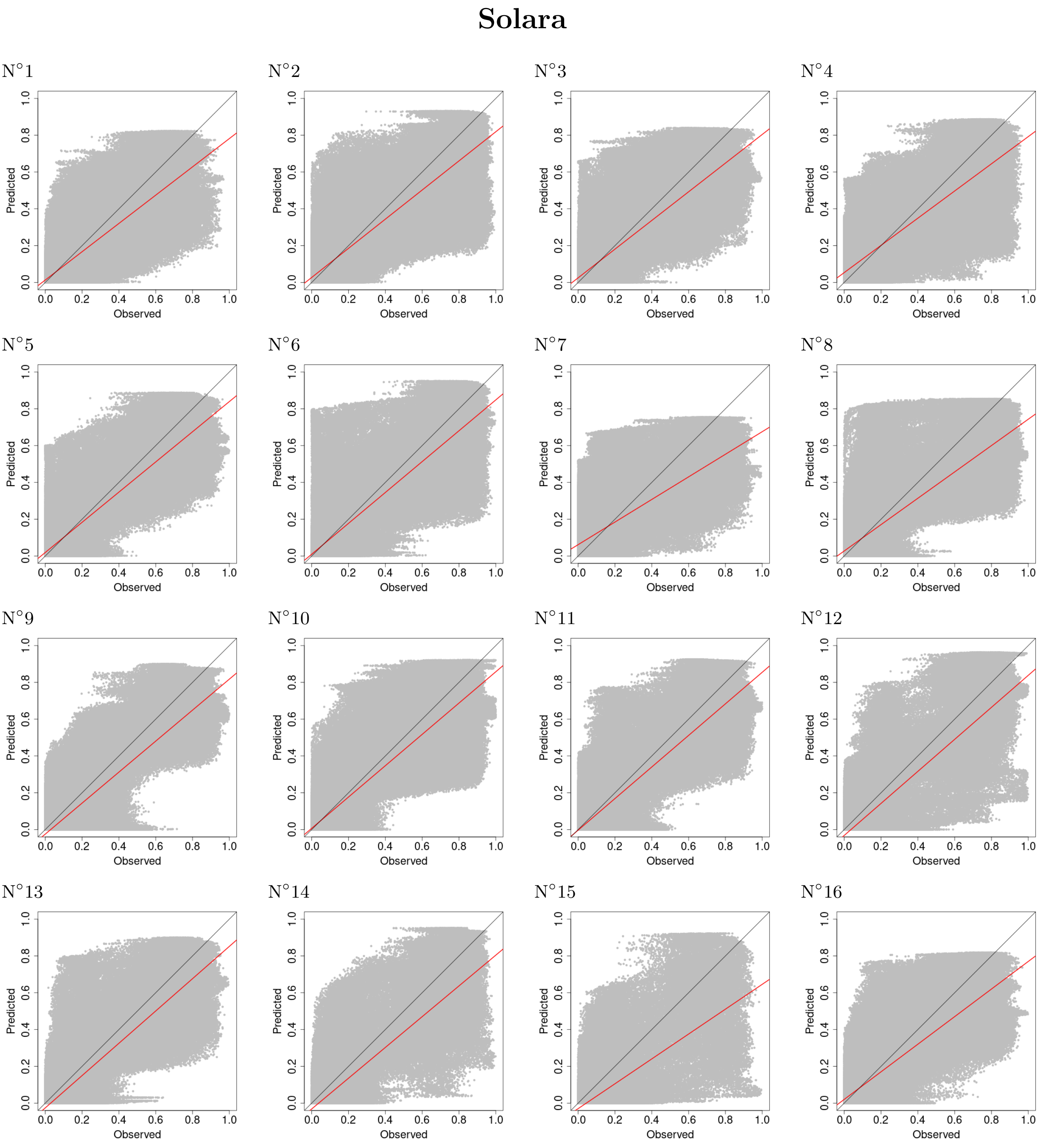


Figure AI: Comparison of infection probability predicted by the reaction-diffusion models against the values of the probability images for the Solara cultivar. The black line is the first bisector that indicates a perfect agreement between values while the red line is the estimated linear relationship between prediction and observation.

### James

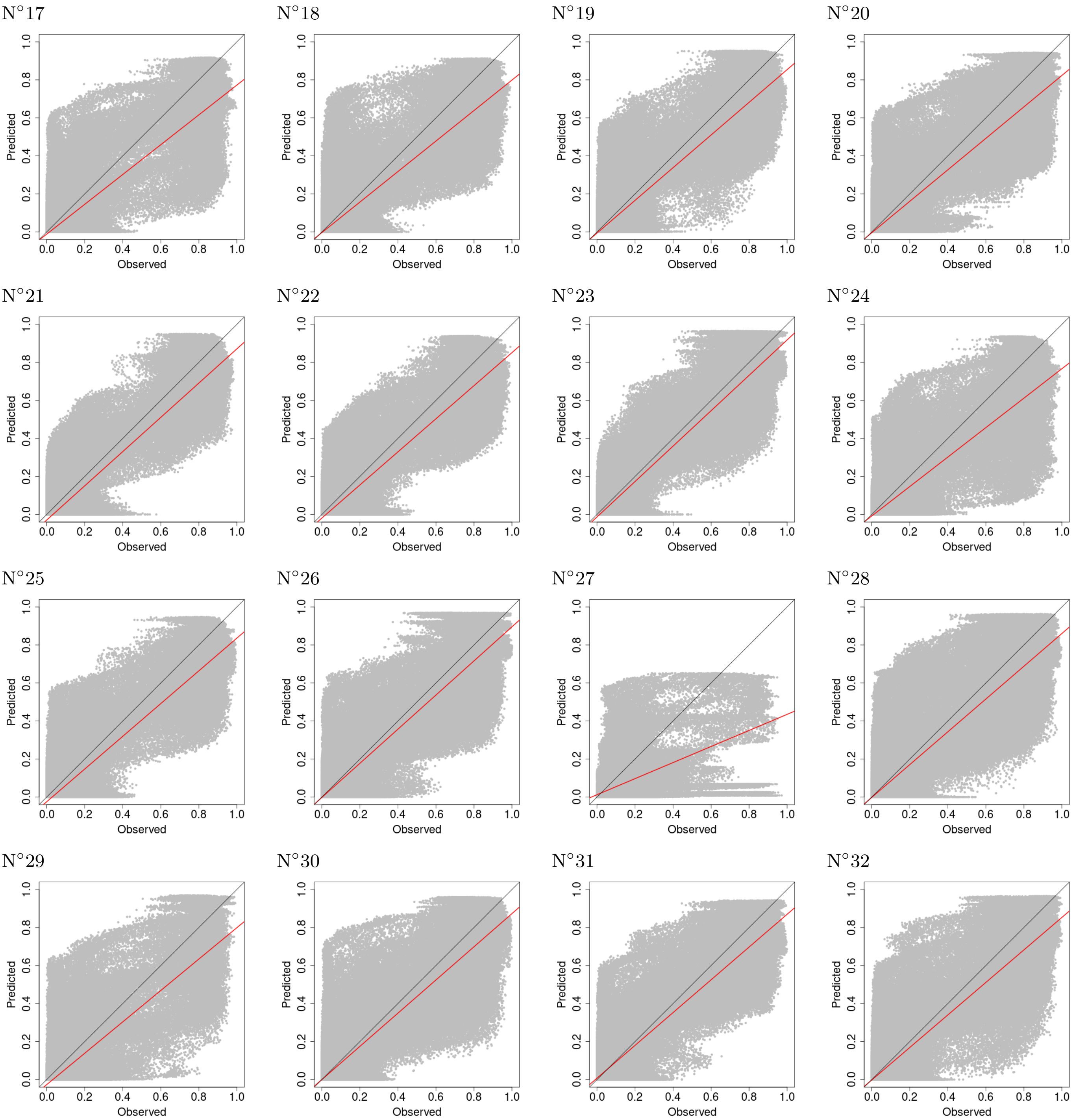


Figure AJ: Comparison of infection probability predicted by the reaction-diffusion models against the values of the probability images for the James cultivar. The black line is the first bisector that indicates a perfect agreement between values while the red line is the estimated linear relationship between prediction and observation.
