## Appendix S2 for "Imaging with spatio-temporal modelling to characterize the dynamics of plant-pathogen lesions"

### S2 Assessment of classifiers quality

Predictions of trained Random Forest classifiers were tested against 31 annotated images obtained from Dutt et al. [1] on the susceptible cultivar Solara and the partially resistant germplasm line DP instead of the cultivar James used in our study. As Dutt et al. [1] did not monitor inoculated stipules 5 days after inoculation we were not able to compare predictions of the classifier for this date. We considered two metrics to compare supervised predictions and ground truth: the balanced accuracy that is the average of sensitivities and specificities, and the Cohen's $\kappa$ that measures the inter-rater reliability in a classification problem. For both metrics the best and worst values are respectively 1 and 0. All the tested classifiers showed a good ability to predict pixel classes with a balanced accuracy ranges from 0.85 to 0.95 and a Cohen's $\kappa$ between 0.77 and 0.96 showing good agreements between supervised segmentation and human annotation (Table A).

| Classifier | Balanced accuracy | Cohen's $\kappa$ |
|---|---|---|
| Day 3 | 0.91 | 0.96 |
| Day 4 | 0.95 | 0.95 |
| Day 6 | 0.85 | 0.77 |
| Day 7 | 0.89 | 0.82 |

Table A: Assessment of classifiers quality on ground truth data with balanced accuracy and Cohen's $\kappa$ metrics.
