## Appendix S1 for "Imaging with spatio-temporal modelling to characterize the dynamics of plant-pathogen lesions"

### S1 Experimental protocol

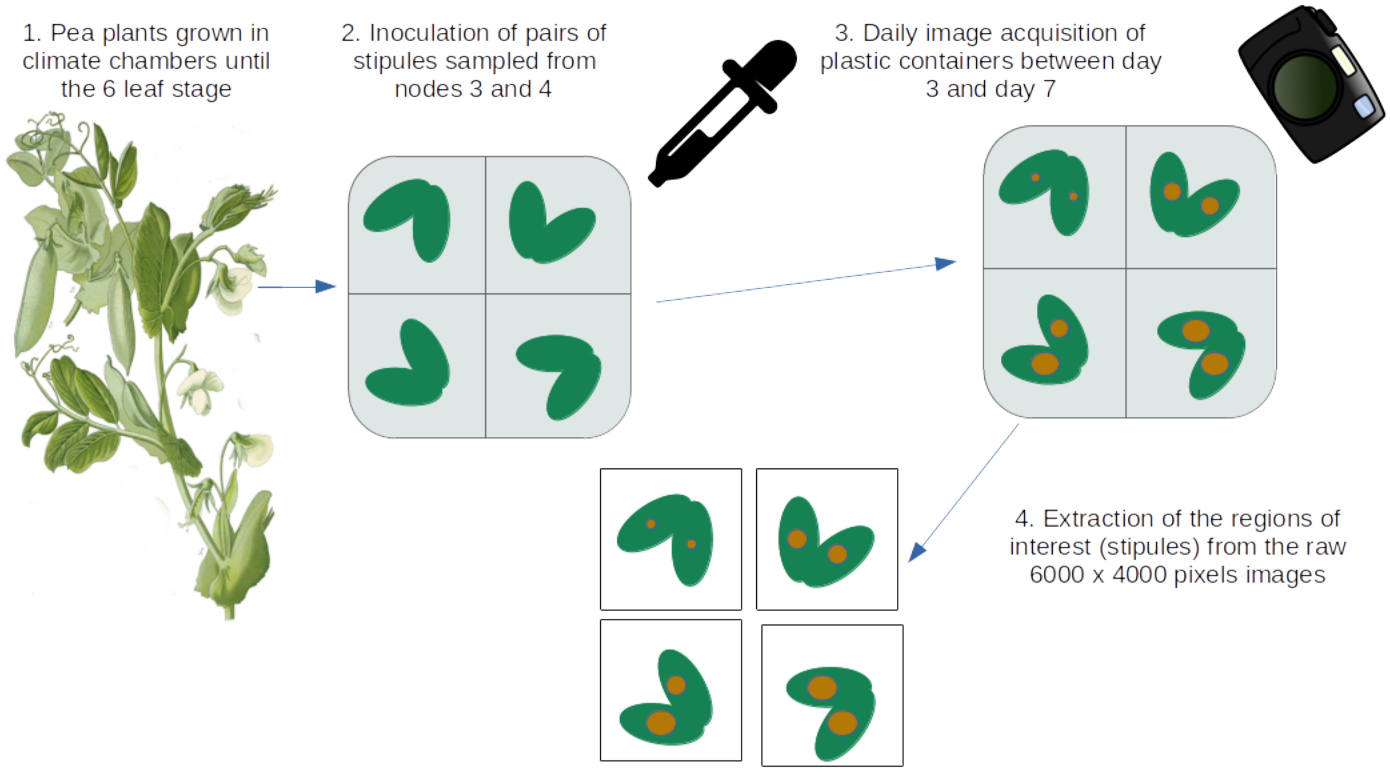


Figure A: Schematic representation of the experimental protocol. The figure was made in LibreOffice impress using open-source cliparts (https://openclipart.org/detail/190961/color-picker for the pipette, https://openclipart.org/detail/203965/camera-digital for the camera) and figure (https://www.biodiversitylibrary.org/page/4321350 for the pea plant).
